# Multilayered Transcriptomic Reprogramming and Spliceosomal Divergence Shape *Bipolaris sorokiniana* Pathogenicity

**DOI:** 10.1101/2025.09.09.675095

**Authors:** Bhakti R. Dayama, Anand Kumar Shukla, Kirtikumar R. Kondhare, Narendra Kadoo

## Abstract

*Bipolaris sorokiniana*, the causative agent of spot blotch of wheat, significantly limits wheat productivity. Despite the pathogen’s widespread impact, the *in planta* molecular mechanisms underpinning its virulence remain poorly characterized. We performed a comprehensive RNA-sequencing analysis of *B. sorokiniana* during attack on spot-blotch-resistant and susceptible wheat genotypes, integrating differentially expressed genes (DEGs), alternative splicing (AS) events, and identification of long non-coding RNAs (lncRNAs). The pathogen exhibited extensive host-genotype-dependent transcriptomic reprogramming, with 128 pathogen genes upregulated in the susceptible host. These genes were associated with ribosome biogenesis, RNA processing, and primary metabolic functions, supporting aggressive colonization. In contrast, pathogen attack on the resistant genotype triggered the upregulation of 58 pathogen genes associated with stress-responsive pathways, including sphingolipid and ceramide metabolism. This suggests a shift toward defensive metabolic reprogramming in the resistant host that restricted its proliferation. Our investigation uncovered five classes of AS events and 14 differentially expressed lncRNAs, revealing substantial post-transcriptional complexity. Notably, a subunit of the H/ACA small nucleolar ribonucleoprotein (snoRNP) complex emerged as a rare “triple-hit” candidate simultaneously identified as a DEG, differentially alternatively spliced gene, and target of a differentially expressed lncRNA, highlighting its potential as a central regulatory node in host-responsive stress adaptation. This study reveals a multilayered regulatory landscape involving transcriptional plasticity, alternative splicing, and lncRNA-mediated control, enabling *B. sorokiniana* to fine-tune its infection strategy in response to host resistance. This work advances the understanding of fungal pathogenesis and identifies molecular vulnerabilities that could be exploited for targeted, host-specific disease control.

## 1. Introduction

Fungal pathogens employ sophisticated strategies to colonize their hosts and cause diseases (Brown 2023). Among them, *Bipolaris sorokiniana*, the causal agent of spot blotch of wheat, is a highly adaptable hemibiotrophic fungus posing a significant threat to global wheat production, especially in warm and humid regions worldwide. It can infect multiple plant tissues, including leaves, roots, and grains. *B. sorokiniana* infection results in substantial yield losses and grain quality deterioration, particularly in susceptible cultivars (Kumar et al. 2022). On average, it causes an annual yield reduction of approximately 17%, with losses in individual fields ranging from 10% to 50%, and in extreme epidemic conditions, up to 100%, depending on the level of cultivar resistance and prevailing environmental conditions (Shukla et al. 2025). Despite its widespread occurrence and agricultural impact, the molecular basis of its pathogenic lifestyle remains underexplored, especially under natural host infection conditions.

Like other hemibiotrophic pathogens, *B. sorokiniana* initiates infection with a biotrophic phase, during which it suppresses host resistance to establish itself. After the pathogen sufficiently proliferates in the host, it switches to the necrotrophic phase marked by the production of toxins or cell wall-degrading enzymes, which results in extensive host tissue degradation. The pathogen employs a diverse arsenal of enzymes, effectors, and toxins to cause disease. These include host-selective toxins and protein effectors, such as *ToxA*, acquired via horizontal gene transfer, and *BsToxA, CsSp1*, and *BsCE66*, all of which contribute to host specificity and tissue colonization (Zhang et al. 2022; Kaladhar et al. 2023; Manan et al. 2023). *B. sorokiniana* also secretes an array of cell wall–degrading enzymes and produces non-host-specific phytotoxins like prehelminthosporol (PHL), with higher production linked to increased virulence (Kumar et al. 2022; Aggarwal et al. 2022). Early infection stages rely on conidial adhesion, mediated by extracellular matrix (ECM) glycoproteins that remodel dynamically during fungal development (Apoga et al. 2001). Furthermore, high genetic diversity driven by parasexuality and heterokaryosis facilitates the emergence of more virulent strains (Basak et al. 2024). Secondary metabolite-associated genes, such as *VHv1* in barley, underscore the importance of specialized biosynthetic pathways in its pathogenic strategy (Zhong et al. 2002). While genome sequencing efforts and *in vitro* transcriptome studies have catalogued many of these candidate virulence-associated genes (Somani et al. 2019; Yadav et al. 2023; Basak et al. 2024), insights into their dynamic regulation within the host context remain limited. Moreover, the transcriptional plasticity that enables *B. sorokiniana* to thrive across contrasting host genotypes, resistant versus susceptible, has not been comprehensively investigated.

In eukaryotic pathogens, transcriptional reprogramming is often complemented by post-transcriptional regulation, including alternative splicing (AS) and long non-coding RNAs (lncRNAs), which enable rapid and fine-tuned adaptation to environmental stress and host resistance (Grützmann et al. 2014; Gonzalez-Hilarion et al. 2016; Kalem and Panepinto 2022). AS enables the generation of multiple transcript isoforms from a single gene, thereby increasing proteomic diversity and functional complexity. In fungal pathogens, AS has been linked to the regulation of genes involved in signal transduction, protein transport, and enzymatic activity during infection (Cheng et al. 2022). Emerging evidence from fungal pathosystems such as *Sclerotinia sclerotiorum, Fusarium graminearum*, and *Magnaporthe oryzae* suggests that post-transcriptional regulatory mechanisms, especially AS and lncRNAs, play critical roles in modulating virulence, stress responses, and cellular reprogramming during infection (Ibrahim et al. 2021; Jeon et al. 2022; Choi et al. 2022; Fu et al. 2024). Recent studies have shown that fungal pathogens can actively reprogram their own RNA splicing machinery to fine-tune gene expression in response to host-derived cues, thereby enhancing their pathogenic potential. In *M. oryzae*, deletion of the serine/arginine-rich splicing factor MoSrp1 led to thousands of mis-spliced transcripts and severely reduced virulence, demonstrating a direct link between AS and pathogenicity. Similarly, splicing regulators, FgSRP1 and FgSRP2, have been shown to be essential for infection in *F. graminearum* (Zhang et al. 2017, 2020). Similarly, over 130 splicing-associated genes in *S. sclerotiorum* were shown to be dynamically expressed during early host invasion, likely as a mechanism to evade plant defenses (Cheng et al. 2022). Additionally, the glycine-rich protein MoGrp1 in *M. oryzae* modulates the splicing of morphogenesis genes necessary for pathogenicity (Gao et al. 2019). These findings suggest that manipulation of RNA splicing is a conserved and critical virulence strategy across diverse fungal pathogens. However, studies on AS in phytopathogenic fungi remain limited, and no data exist for *B. sorokiniana*, particularly during *in planta* infection. Understanding how this pathogen modulates splicing could reveal important regulatory mechanisms involved in host adaptation and pathogenicity, which could be targeted for effective pathogen control.

Similarly, lncRNAs, typically defined as transcripts over 200 nucleotides without protein-coding potential, have recently gained recognition for their roles in regulating gene expression, chromatin remodeling, and RNA processing (Wang and Folimonova 2023). While extensively studied in plants and animals, the identification and functional analysis of lncRNAs in filamentous fungi are limited. LncRNAs have been implicated in regulating plant responses to *Phytophthora infestans, Puccinia graminis*, and *F. graminearum* (Li et al. 2021; Wang and Folimonova 2023). For instance, in *F. graminearum*, the lncRNA lncRsp1 regulates the expression of the virulence-related putative sugar transporter *Fgsp1*, influencing pathogenicity on wheat (Wang et al. 2022), while in the human pathogen *Cryptococcus neoformans*, lncRNAs such as DINOR (DNA damage-inducible non-coding RNA) and RZE1 are required for full virulence and the regulation of infection-related morphogenesis (Kalem and Panepinto 2022). Additionally, in *Ustilago maydis*, antisense lncRNAs expressed during infection have been shown to be essential for pathogenicity; while in *Verticillium dahliae*, specific lncRNAs modulate the expression of cell-wall-degrading enzymes, impacting fungal virulence on cotton (Donaldson and Saville 2013; Choi et al. 2022). A few recent reports in model fungi and human pathogens have uncovered lncRNAs involved in stress responses, pathogenesis, and developmental transitions, yet their roles in fungal–plant interactions remain unknown (Mattick et al. 2023; Wang and Folimonova 2023). Given their regulatory versatility, elucidating the functions of lncRNAs in *B. sorokiniana* could provide valuable insights into the molecular strategies underlying its adaptive infection processes.

In this context, the present study aimed to comprehensively characterize the transcriptional and post-transcriptional landscape of *B. sorokiniana* during infection. By integrating differential gene expression analyses, alternative splicing processes, and lncRNA profiles, we uncovered the molecular mechanisms underpinning pathogen adaptation, virulence, and interaction with the host. This study provides new insights into the regulatory complexity of fungal infection strategies and establishes a foundational resource for identifying novel targets for disease management in cereal crops. Ultimately, the findings of this study will advance the understanding of how *B. sorokiniana* orchestrates its infection strategy for successful host infection and disease development.

## 2. Materials and methods

### 2.1 Plant infection and sample collection

Seedlings of wheat genotypes Chirya3 (spot blotch resistant) and DDK1025 (spot blotch susceptible) were grown under greenhouse conditions and spray-inoculated with spore suspension of *B. sorokiniana* (Dayama et al. 2025). The study showed that the 4 days post-inoculation (dpi) time-point marks a major shift in spot blotch infection in wheat. This time point corresponds to active pathogen growth and host-pathogen interaction. Hence, we selected the 4 dpi time-point to capture fungal virulence-associated gene expression, AS events, and lncRNA profiles during pathogen colonization. We utilized the RNA-sequencing (RNA-seq) data from pathogen-inoculated leaf tissues of Chirya3 and DDK1025 wheat genotypes, which were deposited in the NCBI Sequence Read Archive (SRA) with accession number PRJNA1090403 (Dayama et al. 2025).

### 2.2 RNA-seq data quality control

Quality assessment of the raw sequencing reads was performed using FastQC (Andrews 2010). Key metrics, such as per-base sequence quality, per-sequence quality scores, per-base sequence content, and GC content distribution, were thoroughly examined to ensure data reliability. Adapter sequences, particularly the universal Illumina adapter (AGATCGGAAGAGC), were removed using Trim Galore (v0.6.10) (Krueger 2015). Additionally, reads shorter than 20 nt were discarded. To further improve data quality, low-quality bases were trimmed from the ends of reads, retaining only those with a Phred quality score ≥ 20 (Basak et al. 2024).

### 2.3 Extraction of fungal reads from the dual RNA-Seq data

To extract fungal-specific reads, the reference genomes of *B. sorokiniana* (ND90PR; GCF_000338995.1) and *Triticum aestivum* (bread wheat; GCF_018294505.1) were indexed using HISAT2 (v2.2.1) (Kim et al. 2019). For *B. sorokiniana*, splice-site and exon information were extracted from the corresponding gene annotation file for splice-aware alignment. Trimmed reads were initially mapped to the *B. sorokiniana* ND90PR genome, and the resulting mapped reads were extracted and converted back into paired-end FASTQ format (R1 and R2). These reads were then aligned to the wheat genome to remove potential host-derived sequences. Reads that remained unmapped in this step, which were presumed to be fungal, were remapped to the *B. sorokiniana* genome to refine the fungal read dataset. Finally, Kraken 2 (Wood et al. 2019) was employed to confirm the taxonomic identity of the reads, ensuring their origin from *B. sorokiniana* ND90PR.

### 2.4 Transcriptome assembly and quantification

StringTie (Pertea et al. 2015) was used to quantify gene and transcript expression levels from the fungal reads in each sample. The assembled transcripts were initially merged across all samples using the “merge” option to generate a unified transcriptome reference. This merged reference was used to estimate transcript abundance and generate a count matrix. For this, StringTie was run with the “-e” option to guide the quantification based on the merged transcript annotation. The gffcompare program (Pertea and Pertea 2020) was used to annotate the assembled transcripts. To quantify expression levels, read count matrices at both gene and transcript levels were generated from StringTie output using the *prepDE*.*py3* script (https://ccb.jhu.edu/software/stringtie/dl/prepDE.py), a Python utility provided by StringTie for downstream differential expression analysis. The resulting count matrices were formatted to be compatible with downstream differential expression analysis using DESeq2 (version 1.42.0) (Love et al. 2014) and edgeR (version 4.0.16) (Robinson et al. 2010), the two widely used Bioconductor packages (Pertea et al. 2016).

### 2.5 Differential gene expression and functional analysis

Differential gene expression analysis was carried out using DESeq2 and edgeR. Genes exhibiting a log_2_ fold change (log_2_FC) of > ±1 and a p-value of ≤0.05 were considered significantly differentially expressed and selected for further analysis. A volcano plot was generated using the EnhancedVolcano package (version 1.20.0) (https://github.com/kevinblighe/EnhancedVolcano) to visualize the differentially expressed genes (DEGs). The combined set of DEGs identified by both tools was used for downstream functional annotation. Gene Ontology (GO) enrichment analysis was performed using ShinyGO v0.82 (Ge et al. 2020), with *B. sorokiniana* ND90Pr as the background reference. Enriched GO terms were filtered based on false discovery rate (FDR), and pathways were ranked by fold enrichment. The top enriched biological pathways among upregulated and downregulated genes in both susceptible and resistant wheat genotypes were visualized. A hierarchical clustering tree was generated to represent the correlation among significantly enriched pathways.

For functional annotation, OmicsBox version 3.4 was used (https://www.biobam.com/omicsbox). First, BLASTp (Altschul et al. 1990) was performed against the NCBI non-redundant (nr) protein database (version 5) with a default e-value threshold of 1e^-3^ to identify putative homologs. InterProScan 5.74-105.0 (Blum et al. 2021) was run to retrieve protein domain and family information. GO Mapping (Gotz et al. 2008) was then executed to associate BLAST hits with Gene Ontology terms, followed by GO Annotation using default parameters. Combined GO annotation was performed to summarize GO terms under biological process, cellular component, and molecular function categories. KEGG pathway analysis was also conducted (Kanehisa and Goto 2000), and annotated pathways were grouped into four main categories: (i) Genetic Information Processing, (ii) Cellular Processes, (iii) Metabolism, and (iv) Environmental Information Processing.

To explore potential pathogenicity-related functions, protein sequences derived from the DEGs were queried against the Pathogen-Host Interaction database (PHI-base v4.18) (Urban et al. 2025) using BLASTp. Only high-confidence matches with a minimum sequence identity of 50% and an e-value < 1e^-5^ were retained for further analysis. Carbohydrate-active enzymes (CAZymes) were identified using the dbCAN3 server (https://bcb.unl.edu/dbCAN2/index.php) (Zheng et al. 2023) with DIAMOND (e-value < 1e^−102^) and classified into six major families: Glycoside Hydrolases (GHs), Glycosyltransferases (GTs), Polysaccharide Lyases (PLs), Carbohydrate Esterases (CEs), Auxiliary Activities (AAs), and Carbohydrate-Binding Modules (CBMs). Additionally, effector proteins were predicted from the DEG set using a sequential pipeline (Basak et al. 2024). Initially, the presence of N-terminal signal peptides was assessed using SignalP 6.0 (Teufel et al. 2022), followed by subcellular localization prediction using TargetP 2.0 (Emanuelsson et al. 2000). Transmembrane domains were screened using TMHMM 2.0 (https://services.healthtech.dtu.dk/services/TMHMM-2.0/) (Krogh et al. 2001), and final effector candidates were identified using EffectorP 3.0 (Sperschneider and Dodds 2022).

### 2.6 Network analysis of DEGs in resistant and susceptible wheat genotypes

Network analysis of DEGs was performed separately for upregulated and downregulated genes in both resistant and susceptible wheat genotypes using the STRING database (version 12.0) (Szklarczyk et al. 2023). Protein sequences of the DEGs were mapped to STRING with a medium confidence interaction score threshold of 0.40 to construct protein–protein interaction (PPI) networks. These networks were then examined to identify clusters of interacting proteins, representing potential functional modules. Functional enrichment analysis of the network components was conducted to determine the biological processes associated with the interacting genes. The results were visualized using bubble plots generated using the SRplot tool (Tang et al. 2023), highlighting key functional categories enriched within the DEG networks across different host responses.

### 2.7 Identification of differentially alternative splicing (DAS) events

To explore post-transcriptional regulation in *B. sorokiniana* during its interaction with resistant and susceptible wheat hosts, we performed alternative splicing (AS) analysis using SUPPA2 (Trincado et al. 2018), a robust tool for isoform-level splicing quantification. Splicing events were generated from the merged transcript annotation, and transcripts per million (TPM) values were extracted using StringTie and prepDE.py3. Each event’s percentage or proportion spliced-in (PSI) values were calculated using psiPerEvent, followed by differential splicing analysis (diffSplice) between resistant and susceptible wheat genotypes. Significant DAS events were filtered based on ΔPSI > 0.2 and p-value < 0.5. The genes corresponding to significant DAS events were subjected to GO enrichment using ShinyGO (Ge et al. 2020), and further annotated using PHI-base as described for DEGs.

### 2.8 Identification and characterization of lncRNAs

The lncRNAs were identified from the assembled transcriptome through a series of stringent filtering steps (Yan et al. 2020; Zhou et al. 2022; Shi et al. 2025). Initially, transcripts longer than 200 nucleotides (Lee and Kikyo 2012) and containing at least two exons were selected to obtain multi-exonic candidates. Novel transcripts were then extracted based on specific StringTie class codes (u, p, m, n, o, j, x, i). A minimum expression threshold of TPM ≥ 1 in at least one condition was applied, resulting in a set of high-confidence novel lncRNAs. These candidates were further screened against the Rfam (Ontiveros-Palacios et al. 2025) and Pfam databases (Mistry et al. 2021) using an e-value cutoff of <0.001. Additionally, transcripts containing excessive ambiguous bases (‘N’) were removed. To assess coding potential, the remaining transcripts were analyzed using CPC2 (Kang et al. 2017) and CPAT (Wang et al. 2013), with a coding probability threshold of <0.5, confirming their non-coding nature. Differential expression analysis of lncRNAs between resistant and susceptible wheat genotypes was performed using DESeq2 and edgeR, applying a significance threshold of p-value < 0.2 and log_2_ fold change ≥ ±2. The union of differentially expressed lncRNAs (DElncRNAs) from both tools was considered for downstream analysis. Subcellular localization of the lncRNAs was predicted using the iLoc-LncRNA tool (http://lin-group.cn/server/iLoc-LncRNA/pre.php) (Su et al. 2018), enabling functional classification based on predicted cellular compartments. Further, genomic classification of lncRNAs relative to nearby protein-coding genes was conducted using class codes from the StringTie annotation (Pertea et al. 2015).

### 2.9 Integration network of DEGs, DAS genes, and lncRNAs

An integrated network of DEGs, DAS genes, and DElncRNAs was constructed to explore potential regulatory relationships. Cis-targets of DElncRNAs were identified by locating protein-coding DEGs and DAS genes within a ±100 kb genomic window using BEDTools 2.31.1 (Zhou et al. 2022; Xia et al. 2025), while trans-targets were predicted based on Spearman correlation analysis (r ≥ 0.9, p < 0.05) between DElncRNAs and DEGs/DAGs (Augustino et al. 2020). The resulting interaction networks were visualized and analyzed using Cytoscape (version 3.10.3) (Shannon et al. 2003) to reveal potential regulatory modules and functional associations among transcripts.

## 3. Results

### 3.1 RNA sequencing and extraction of *B. sorokiniana* transcripts

Our previous study (Dayama et al. 2025) identified 4 dpi as a critical stage marked by visible disease progression and active host-pathogen interaction. Hence, we focused this study on the 4 dpi stage. This time point was strategically chosen to capture the dynamic transcriptional landscape of *B. sorokiniana*, enabling investigation of its virulence-associated gene expression, AS events, and lncRNAs during early stages of host colonization. RNA-seq generated approximately 589 million raw reads across the six samples, with individual libraries yielding between 60 and 137 million reads (Table S1). The GC content ranged from 45–51% and the percentage of high-quality bases (Q ≥ 20) exceeded 94% in all samples, ensuring reliable data for subsequent analysis. The RNA-seq analysis workflow is presented in Fig. 1a. Given the dual nature of RNA-seq from infected leaf tissue, a multi-step alignment approach was applied to enrich the pathogen-derived sequence reads. As expected, a higher proportion of *B. sorokiniana* reads was recovered from susceptible wheat samples (0.07%; 74,714 of 10,60,45,134 reads) as compared to the samples from the resistant wheat genotype (0.04%; 83,499 of 18,79,48,163 reads) (Table S2). This difference mirrors phenotypic observations, where susceptible plants displayed more severe symptoms and higher fungal biomass than resistant ones. Kraken2 classification confirmed that over 99.5% of the extracted reads in all samples were confidently assigned to *B. sorokiniana*. This high-confidence fungal transcript set was used for downstream analyses.

**Fig. 1.**
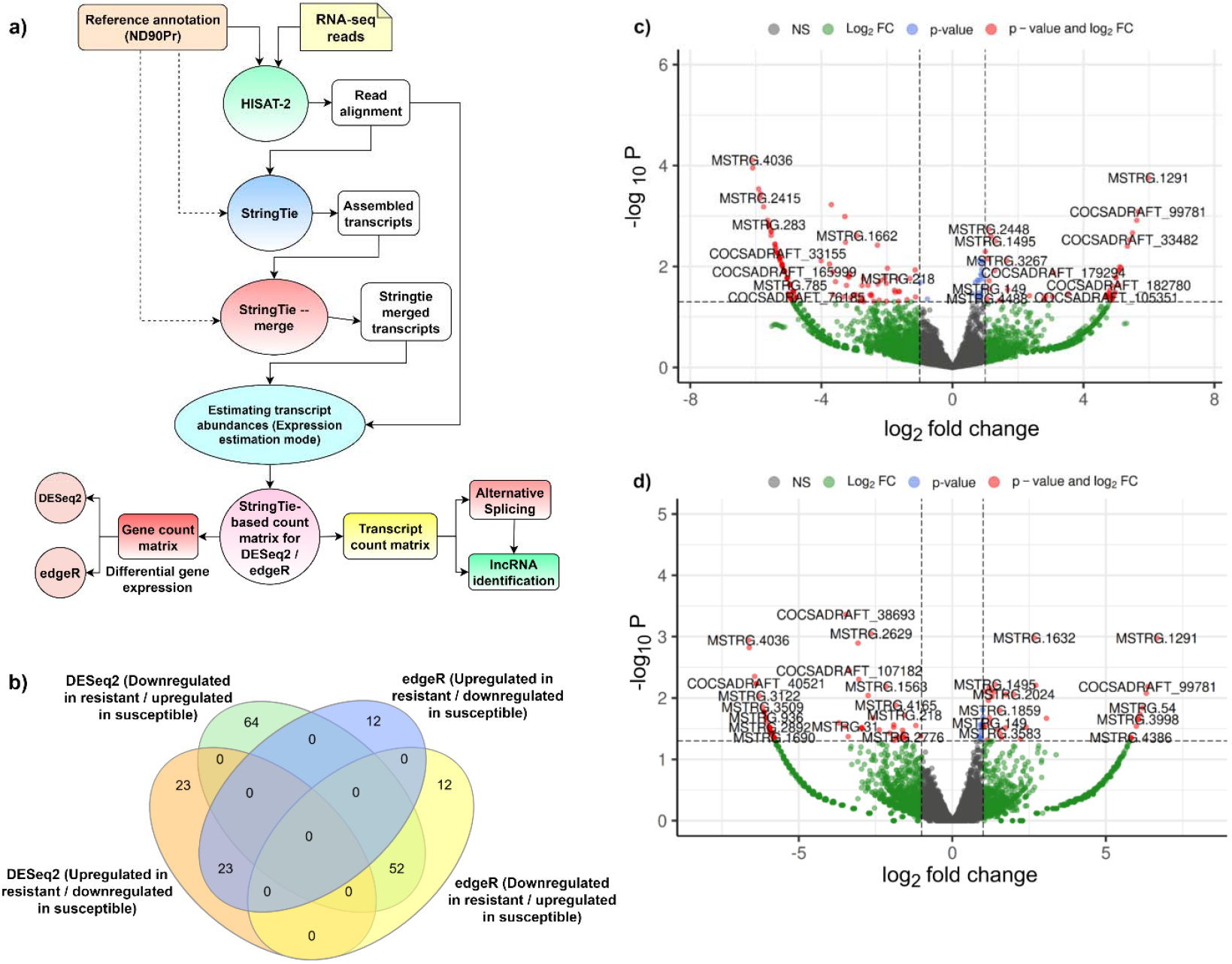
Workflow and differential expression analysis of *Bipolaris sorokiniana* transcriptome in resistant and susceptible wheat genotypes. a) Overview of the RNA-seq data processing and analysis pipeline. b) Venn diagram illustrating the overlap of DEGs identified by DESeq2 and edgeR methods, c) Volcano plot displaying DEGs determined by DESeq2 analysis. d) The volcano plot displays DEGs determined by the EdgeR analysis. Significant DEGs were defined by p-value < 0.05 and absolute log_2_ fold change ±1. Genes upregulated in the resistant genotype and downregulated in the susceptible genotype are labeled as UpR/DownS, while those downregulated in the resistant and upregulated in the susceptible are labeled as DownR/UpS.

### 3.2 Differential gene expression in *Bipolaris sorokiniana* in susceptible and resistant hosts

To decipher the transcriptional differences in *B. sorokiniana* infecting susceptible (SI) and resistant (RI) wheat genotypes, differential gene expression analysis was conducted using DESeq2 and edgeR pipelines. Using DESeq2, 162 DEGs were detected, with 46 genes upregulated in RG and downregulated in SG (UpR/DownS), and 116 downregulated in RG and upregulated in SG (DownR/UpS) (Tables S3-S5). Parallel analysis with edgeR revealed 99 DEGs, including 35 UpR/DownS and 64 DownR/UpS genes (Tables S3-S8). A total of 186 DEGs were identified, with a subset showing consistent regulation patterns across both methods, in which 58 were UpR/DownS and 128 DownR/UpS (Fig. 1b). Volcano plots revealed multiple fungal transcripts significantly upregulated in the susceptible genotype, including putative virulence-associated genes, such as *MSTRG*.*4377* (*epoxide hydrolase*), *MSTRG*.*4687* (*FAD Domain containing protein*), *MSTRG*.*936* (*Gar1-domain containing protein*) and *COCSADRAFT_40521* (*glycerol-3-phosphate mitochondrial precursor*) (Fig. 1c-d). In contrast, genes like *MSTRG*.*3639* (putative transporter), *MSTRG*.*4516* (fungal-specific transcription factor domain), and *COCSADRAFT_182780* (MFS transporter), etc., were the upregulated fungal genes in the resistant genotype (Fig. 1c-d). These findings highlight transcriptional reprogramming in *B. sorokiniana* in response to differing host resistance levels.

### 3.3 Functional annotation and GO categorization of *B. sorokiniana* DEGs

To understand the functional relevance of transcriptional changes in *B. sorokiniana* during interaction with RG and SG, GO enrichment analysis was performed using pathogen DEGs combined from DESeq2 and edgeR analysis. The DownR/UpS fungal genes were significantly enriched for processes related to ribosome biogenesis and spliceosomal function, alongside oxidative stress response and carbohydrate metabolism. The GO terms, such as ribonucleoside-diphosphate reductase activity, oxidoreductase activity, and polysaccharide catabolic process, indicated enhanced transcriptional and translational activity, supporting accelerated fungal growth and metabolic demand during successful colonization in the susceptible wheat genotype (Fig. S1a). Conversely, the UpR/DownS genes were enriched for functions associated with cell wall biosynthesis, intracellular transport, and structural maintenance. GO terms such as fungal-type cell wall organization, chitin synthase activity, glucanosyltransferase activity, sphingolipid biosynthesis, and vesicle-mediated transport indicate activation of stress-responsive pathways (Fig. 1b).

Functional annotation of the 128 DownR/UpS and 58 UpR/DownS genes was performed using OmicsBox, incorporating BLAST searches against the NCBI nr database, InterProScan domain analysis, and Gene Ontology (GO) mapping (Fig. S2, Tables S9-S10). Combined GO analysis revealed contrasting molecular responses of *B. sorokiniana* during infection of resistant and susceptible wheat genotypes. In the DownR/UpS category, biological processes, such as organelle organization, metabolic and catabolic processes, transmembrane transport, and ribonucleoprotein complex biogenesis, were abundant, suggesting enhanced fungal cellular activity, molecular transport, and biosynthetic functions that facilitate successful colonization in the susceptible host. Conversely, in the UpR/DownS category, enrichment of small molecule metabolism, methylation, and membrane organization indicated a stress-adaptive metabolic reprogramming of the pathogen in response to the resistant host environment, likely reflecting its efforts to maintain cellular homeostasis and cope with host defense mechanisms. At the molecular function level, elevated hydrolase and oxidoreductase activities in the DownR/UpS group reflected increased oxidative stress handling and proteolytic processes during active infection. In contrast, resistance-specific activation of pathways such as zinc ion binding, ribonucleoside triphosphate phosphatase activity, and methyltransferase activity pointed to roles in signal transduction, transcriptional regulation, and chromatin remodeling under defense-triggered conditions. For cellular components, the fungal-type cell wall, nucleus, and intracellular protein-containing complex were more abundant in the UpR/DownS group, suggesting reinforcement of structural integrity and intracellular regulation during stress. In contrast, enrichment of nuclear protein-containing complexes, nuclear lumen, endomembrane system, and mitochondria in DownR/UpS suggests elevated biosynthetic and energy-generating activities in the susceptible host (Fig. S2).

To further dissect the functional landscape of *B. sorokiniana* genes modulated by host genotype, a deeper exploration of the DEGs (DownR/UpS and UpR/DownS) was carried out using KEGG pathways, PHI-base virulence annotations, CAZy enzyme classification, and effector predictions to pinpoint potential pathogenicity mechanisms (Fig. S3). Differential expression analysis revealed that the DownR/UpS fungal genes were primarily enriched in metabolism (58 genes) and cellular processes (15 genes), indicating active fungal growth and metabolic activity. In contrast, UpR/DownS genes showed fewer enrichments, with metabolism (18 genes) being the most represented, suggesting a stress-adaptive or compensatory response under host resistance (Fig. S3a, Tables S11-12).

Annotation of the fungal DEGs against the PHI-base database revealed distinct shifts in virulence-associated functional categories between resistant and susceptible wheat hosts (Fig. S3b, Table S13). The DownR/UpsS genes were predominantly associated with reduced virulence (11 genes) and unaffected pathogenicity (5 genes), with additional representation in categories such as loss of pathogenicity, lethal loss of pathogenicity, and effector (plant avirulence determinant). This suggests that successful infection in the susceptible host may involve the active expression of genes with known roles in pathogenicity and host interaction. Conversely, the UpR/DownS genes were primarily linked to reduced virulence (7 genes) and unaffected pathogenicity (4 genes), indicating a potential suppression of key virulence factors in response to host resistance. These findings underscore the influence of host genotype on the transcriptional regulation of fungal pathogenicity determinants (Fig. S3b; Table S13).

CAZyme (carbohydrate-active enzymes) annotation was performed to identify enzymes involved in plant cell wall degradation, a key step in fungal penetration and colonization (Fig. S3c, Table S14). Annotation of fungal DEGs against the CAZy database revealed a clear enrichment in the susceptible host background. The DownR/UpS genes included higher numbers of glycoside hydrolases (13 genes), glycosyltransferases (8 genes), and auxiliary activity enzymes (6 genes), indicating enhanced capacity for host cell wall degradation and nutrient acquisition. In contrast, UpR/DownS genes showed markedly fewer CAZyme annotations, with only five genes in glycoside hydrolases and minimal abundance across other CAZyme families. This suggests that the resistant host may suppress fungal enzymatic machinery essential for colonization and pathogenicity (Fig. S3c; Table S14). Similarly, effector analysis identified a subset of DEGs with potential roles in host-pathogen interactions. Notably, four apoplastic effectors and one cytoplasmic/apoplastic effector (MSTRG.1830; Uncharacterized protein) were upregulated in the susceptible host but downregulated in the resistant host, indicating their possible involvement in successful colonization and immune evasion during susceptible infections. No effector candidates were upregulated in the resistant genotype, suggesting that host resistance may suppress effector expression or activity as part of its defense strategy (Fig. S3d; Table S15).

### 3.4 Protein–protein interaction network analysis reveals host-dependent functional specialization

To investigate the functional interplay among the DEGs, protein-protein interaction (PPI) networks were constructed using the STRING database (https://string-db.org). The DownR/UpS gene set formed a highly interconnected network comprising 145 nodes and 329 edges (PPI enrichment *p* < 1.0 × 10^−16^), indicating significant biological connectivity (Fig. 2). Gene Ontology (GO) enrichment of this network revealed a strong association with spliceosomal complex components, RNA splicing, and mRNA processing, suggesting enhanced post-transcriptional regulatory activity during infection of the susceptible host. In contrast, the UpR/DownS gene set formed a smaller but statistically significant network with 73 nodes and 66 edges (*p* = 5.92 × 10^−11^) (Fig. 2a). These genes were predominantly enriched in sphingolipid biosynthesis, ceramide catabolic processes, and sulfate assimilation, pathways associated with membrane remodeling and stress adaptation (Fig. 2b).

**Fig. 2.**
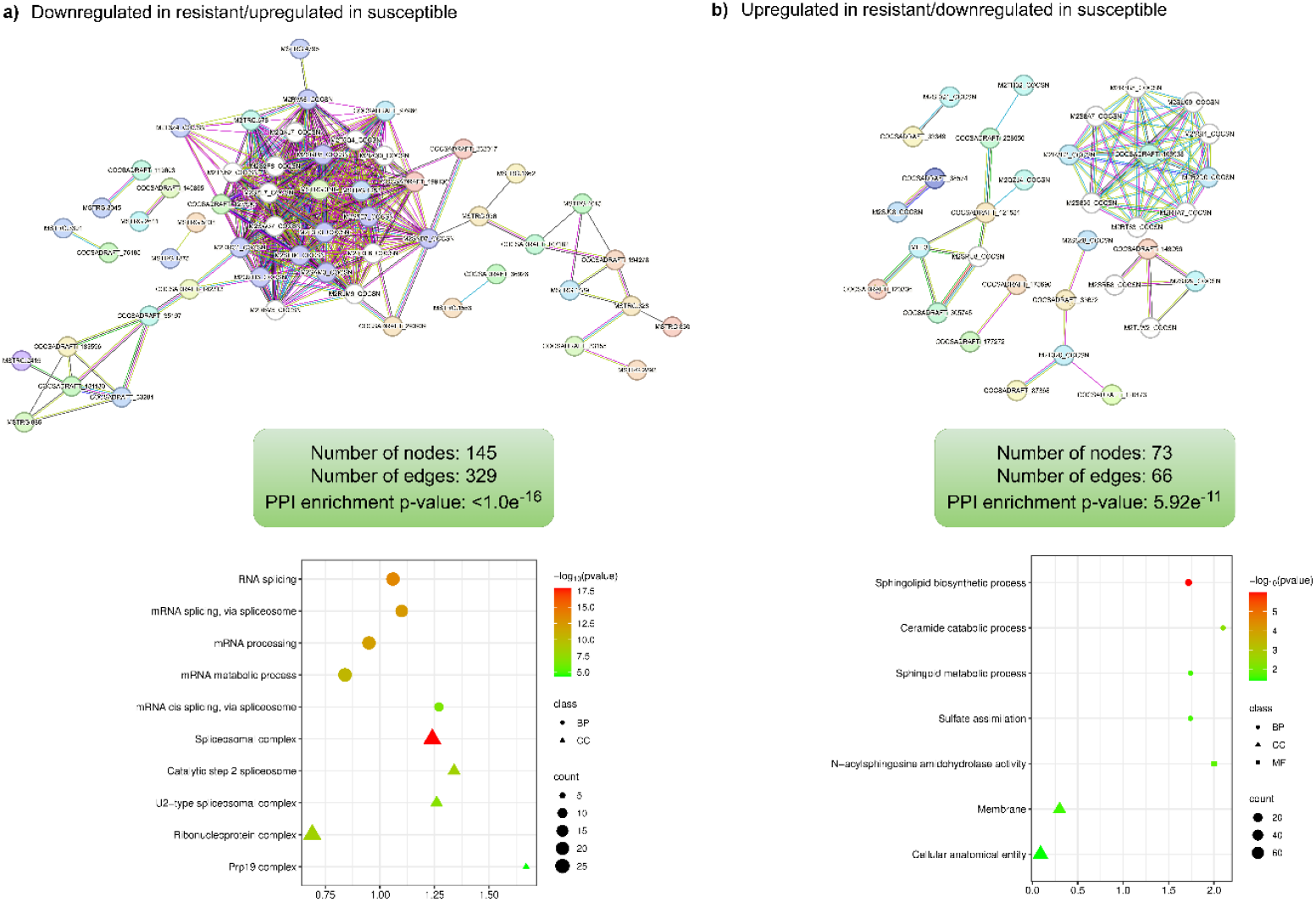
Protein–protein interaction networks and Gene Ontology enrichment analysis of *Bipolaris sorokiniana* DEGs. a) Protein–protein interaction (PPI) network and corresponding Gene Ontology (GO) enrichment for genes downregulated in RG and upregulated in SG (DownR/UpS). b) PPI network and GO enrichment for genes upregulated in RG and downregulated in SG (UpR/DownS). Bubble sizes correspond to gene count, color represents statistical significance (–log10 p-value), and shape indicates GO category: BP (circle), CC (triangle), and MF (square).

### 3.5 Alternative splicing landscape of *Bipolaris sorokiniana* during infection

To explore post-transcriptional regulation in *B. sorokiniana* during its interaction with resistant and susceptible wheat hosts, alternative splicing (AS) analysis was performed using SUPPA2, a robust tool for isoform-level splicing quantification. This study presents the first *in planta* assessment of AS in this fungal pathogen. In our transcriptome datasets, 116 AS events were identified, which included five classical splice types: retained introns (RI), alternative 5⍰ splice sites (A5SS), alternative 3⍰ splice sites (A3SS), alternative first exons (AF), and exon skipping (SE) (Fig. 3a). We found that RI was the predominant splicing event in *B. sorokiniana*, with 52 instances, followed by A5SS (30), A3SS (29), AF (3), and SE (2) (Fig. 3b, Tables S16-S21).

**Fig. 3.**
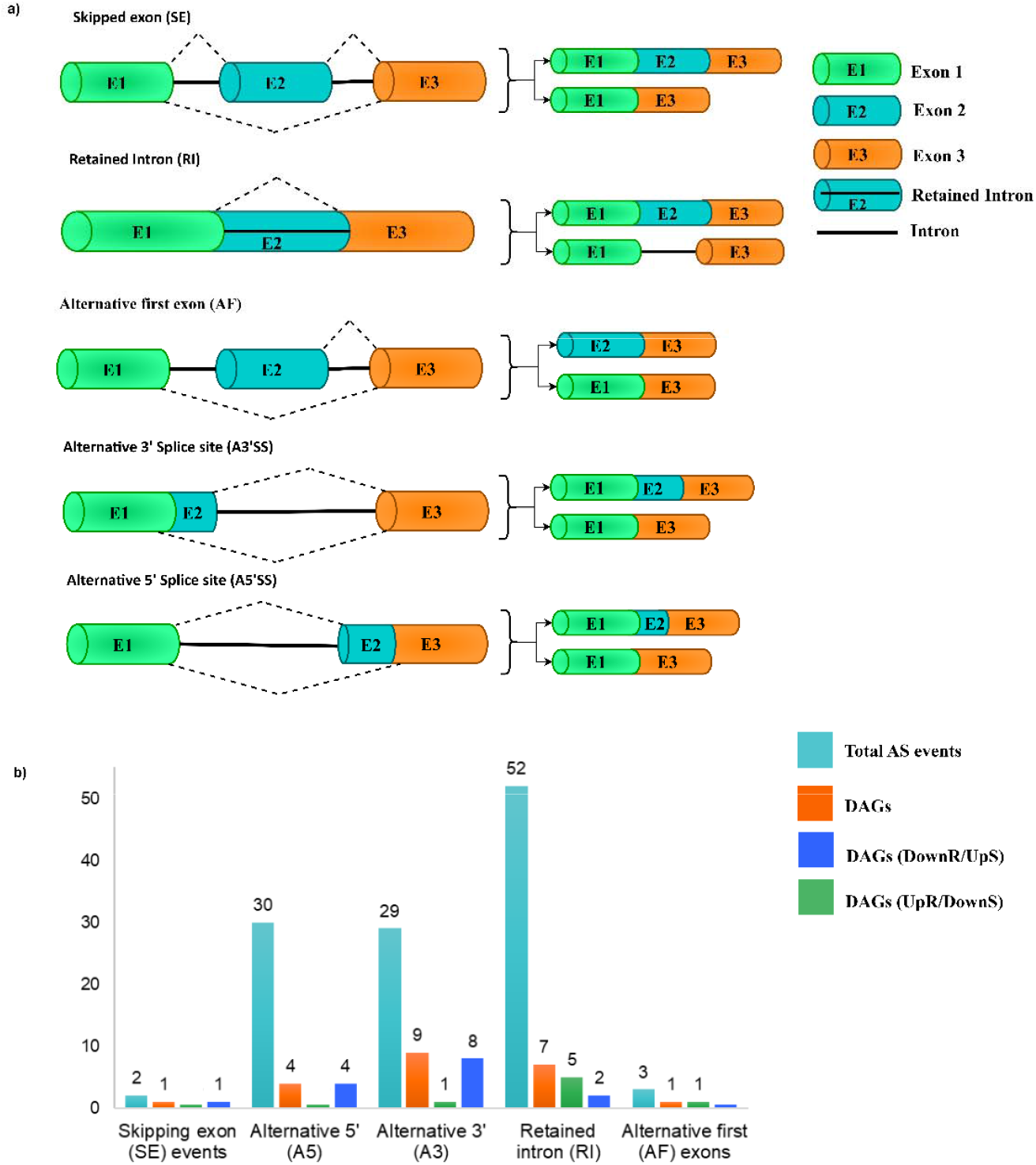
Overview of alternative splicing dynamics in *Bipolaris sorokiniana* during infection of resistant and susceptible wheat genotypes. a) Schematic representation of five major types of alternative splicing (AS) events: Skipped exon (SE), Retained intron (RI), Alternative first exon (AF), Alternative 3’ splice site (A3SS), and Alternative 5’ splice site (A5SS). b) Bar chart showing the number of genes involved in each AS event type across four categories: Total AS genes (turquoise), Differentially alternatively spliced genes (DAGs) (orange), Upregulated in resistant/downregulated in susceptible genotypes (UpR/DownS) (green), and downregulated in resistant/upregulated in susceptible genotypes (DownR/UpS) (blue).

Differential alternative splicing analysis (ΔPSI > 0.2; p-value < 0.5) revealed 22 DAS events, with seven enriched in the resistant host interaction (UpR/DownS) and 15 in the susceptible host (DownR/UpS) (Fig. 4b). The dominance of RI events suggests a potential role for AS in modulating transcript stability or nonsense-mediated decay (NMD) in response to host-imposed stress. The less frequent SE and AF events may reflect a more specialized regulatory mechanism activated under specific host conditions. Several DAS events were detected in genes associated with key biological functions relevant to pathogenicity and cellular adaptation in *B. sorokiniana*. These included *catalase (MSTRG*.*1679), methionine aminopeptidase (MSTRG*.*1971), E3 ubiquitin-protein ligase (MSTRG*.*4464)*, and *plasma membrane ATPase (MSTRG*.*4735)*, each exhibiting intron retention or splice site variation. Additional splicing events affected transcripts, such as *40S ribosomal protein S6* (*MSTRG*.*107*), *translation initiation factor IF2/IF5 (MSTRG*.*4249)*, and *mitochondria-associated protein 19 (MSTRG*.*2223)*, indicating dynamic post-transcriptional regulation of core cellular processes. Together, these results suggest that AS contributes to the functional plasticity of *B. sorokiniana*, enabling it to adapt differentially to resistant and susceptible wheat genotypes.

**Fig. 4.**
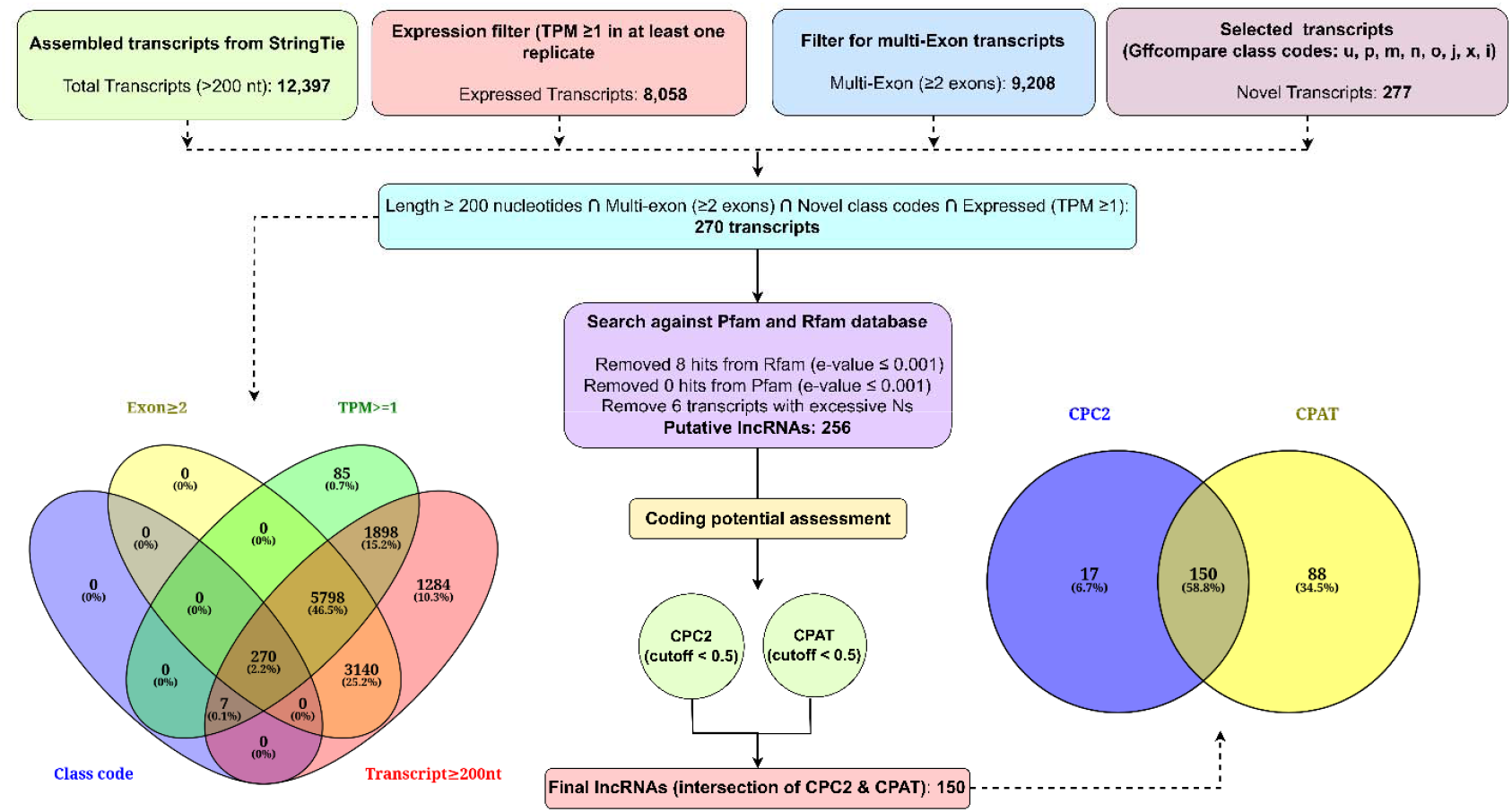
Workflow for identifying high-confidence long non-coding RNAs in *Bipolaris sorokiniana* during wheat infection. Transcripts assembled using StringTie were filtered sequentially based on transcript length (≥200 nt), expression level (TPM ≥1 in at least one replicate), exon number (≥2), and GffCompare-defined novel class codes (u, p, m, n, o, j, x, i). The intersection of these criteria yielded 270 high-confidence candidate long non-coding RNAs (lncRNAs). Further filtering using Rfam and Pfam databases removed 14 transcripts (8 with Rfam hits, 6 with ambiguous nucleotides). The remaining 256 putative lncRNAs were subjected to coding potential prediction using CPC2 and CPAT, with a coding probability cutoff of <0.5. The intersection of both tools identified 150 final lncRNAs. Venn diagrams summarize the transcript filtering and coding potential assessment steps.

GO enrichment analysis of differentially alternatively spliced genes revealed contrasting functional profiles between the UpR/DownS and DownR/UpS (Fig. S4). In the UpR/DownS group, enrichment was observed for RNA splicing and protein metabolism processes, including the U12-type spliceosomal complex, U4 snRNP, SMN-Sm protein complex, and cellular protein metabolic processes. These findings suggest that AS events in the resistant genotype may modulate components of the spliceosome and post-translational machinery, possibly contributing to stress adaptation or fine-tuning of gene expression. Conversely, DAS genes in the DownR/UpS group were enriched for translation initiation factor binding, RNA polymerase II core complex, splice site selection, and protein insertion into the ER membrane. These pathways are linked to enhanced transcriptional and translational activity. This indicates that AS may promote increased expression and processing of proteins involved in active fungal growth and infection in the susceptible genotype. These results suggest a host-dependent splicing regulation, with the interactions in the resistant genotype favoring modulation of spliceosomal machinery and protein turnover. In contrast, the interactions in the susceptible genotype promote splicing variants associated with increased biosynthetic and transcriptional activity.

Annotation of differentially alternatively spliced genes against the PHI-base database identified five candidate genes associated with experimentally validated pathogenicity phenotypes across multiple fungal pathogens. Among these, *MSTRG*.*1679 (catalase)* and *MSTRG*.*4865 (serine/threonine-protein phosphatase)* were associated with unaffected pathogenicity, suggesting that their alternative splicing may not significantly alter virulence under the tested conditions. In contrast, *MSTRG*.*4962 (spermidine spermine synthase)* and *MSTRG*.*385 (serine threonine-protein phosphatase)* were linked to multiple virulence-related phenotypes, including reduced virulence, loss of pathogenicity, and increased virulence. MSTRG.4962 was orthologous to *Fgspe3* from *F. graminearum* and *Spe-Sdh* from *U. maydis*, both associated with impaired virulence upon disruption. MSTRG.385 exhibited homology to genes, such as *SIT4* and *PPG1*, previously implicated in reduced virulence and hypervirulence phenotypes in *Candida albicans* and *F. graminearum*. Additionally, *MSTRG*.*4735 (plasma membrane ATPase)* showed similarity to *Lmpma1* and *ChPMA2*, both linked to loss of pathogenicity, indicating that splicing variation in this gene may influence critical virulence pathways. These findings suggest that AS may modulate key virulence-associated genes in *B. sorokiniana*, potentially impacting its pathogenicity depending on host context and environmental conditions (Table S22).

### 3.6 Identification of long non-coding RNAs (lncRNAs)

We also identified lncRNAs from the *B. sorokiniana* transcriptome captured at 4 dpi from infected wheat tissues to uncover potential regulatory layers contributing to fungal virulence. Starting with 12,397 assembled transcripts (≥200 nucleotides) from StringTie, a stepwise filtering pipeline was employed (Fig. 4) to ensure stringency, novelty, and expression reliability. From this dataset, 8,058 transcripts met the expression criterion (TPM ≥ 1 in at least one replicate), while 9,208 transcripts contained two or more exons. Further reducing the number of transcripts based on GffCompare-defined novel class codes (u, p, m, n, o, j, x, i) led to 277 putatively novel transcripts. The intersection of all four filters, length ≥200 nt, multi-exonic structure, novel class, and expression ≥1 TPM, yielded a final pool of 270 high-confidence candidate lncRNAs. A search against the Rfam and Pfam databases (e-value ≤ 0.001) was performed to remove transcripts with homology to known RNA families. This eliminated eight transcripts with Rfam hits, while Pfam analysis revealed no detectable protein domains. Additionally, six transcripts with excessive ambiguous nucleotides (Ns) were discarded, resulting in 256 putative non-coding transcripts. To further validate their non-coding nature, we used CPC2 and CPAT, which predict coding potential based on sequence features. CPC2 classified 167 transcripts as non-coding, and CPAT identified 238 as non-coding. The overlap of both predictions revealed 150 high-confidence lncRNAs, which we considered as the core set for downstream analysis (Tables S23-S28). This marks the first comprehensive identification of lncRNAs in *B. sorokiniana* during host infection, laying a foundation for understanding their potential roles in fungal adaptation and virulence.

To investigate the regulatory roles of lncRNAs in *B. sorokiniana* during host infection, utilizing DESeq2 and edgeR tools, we performed differential expression analysis of the 150 high-confidence lncRNAs between resistant (RG) and susceptible (SG) wheat genotypes using samples from the 4 dpi time-point. Through this combined approach, we identified 14 differentially expressed lncRNAs (DElncRNAs) (*p*-value < 0.2 and log_2_ fold change ≥ ±2) consistently supported by both methods. Among these, 13 lncRNAs were significantly downregulated in RG and upregulated in SG (DownR/UpS), indicating their potential involvement in enhancing fungal virulence or adaptation within the susceptible host environment (Fig. S5a-b). In contrast, only a single lncRNA, MSTRG.831.2, was of the type UpR/DownS. Among the DownR/UpS lncRNAs, MSTRG.2446.2, MSTRG.4809.1, and MSTRG.936.1 exhibited the highest fold changes and strongest statistical support, indicating their possible roles as key regulatory hubs during host-pathogen interaction. The convergence of results from DESeq2 and edgeR further reinforces the reliability of these findings and highlights a core set of lncRNAs potentially involved in modulating pathogenicity mechanisms in *B. sorokiniana* (Fig. S5c, Tables S29-S30).

Further characterization of the 14 DElncRNAs based on predicted subcellular localization revealed that the majority were localized to the cytoplasm (n = 5) and ribosome (n = 5), followed by the exosome complex (n = 2) and nucleus (n = 2) (Fig. S5d). These localization patterns suggest that most DElncRNAs may function post-transcriptionally, potentially regulating mRNA stability, translation, or ribosome-associated processes, particularly during host infection in the susceptible genotype. In addition, analysis of the genomic context of these DElncRNAs showed a predominance of sense lncRNAs (n = 11), with fewer classified as antisense (n = 2) and intergenic lncRNAs (n = 1) (Fig. S5e). The high proportion of sense-oriented lncRNAs implies that they may act in *cis* mode to modulate the expression or processing of overlapping protein-coding genes. In contrast, antisense and intergenic lncRNAs may function via trans-regulatory mechanisms. Together, these findings provide insight into the possible regulatory roles and functional diversity of DElncRNAs during *B. sorokiniana* infection under different host resistance conditions.

### 3.7 Co-expression network analysis of DEGs, DAGs, and DElncRNAs

To explore regulatory interactions underlying *B. sorokiniana* pathogenicity, we constructed a comprehensive cis- and trans-acting co-expression network integrating DEGs, DAGs, and DElncRNAs during infection of resistant and susceptible wheat genotypes. In the cis-regulatory network, five DElncRNAs were associated with seven target genes (Fig. 5a, Table S31), including three DEGs, two DAGs, and one gene common to both categories. Among these, five targets were UpR/DownS genes, while two exhibited the opposite pattern (DownR/UpS). Notably, MSTRG.936.1 was linked to both DEGs and DAGs and was itself differentially expressed, suggesting a potential multifaceted role in local gene regulation. The trans-regulatory network (Fig. 5b; Table S32) revealed interactions between six DElncRNAs and 39 distant target genes, comprising 38 DEGs and one DAG. Of these, 26 genes were DownR/UpS, implicating these targets in fungal virulence or adaptation in the susceptible host environment. The remaining 13 genes were UpR/DownS, possibly reflecting the desperate attempts by the pathogen to establish itself under host resistance.

**Fig. 5.**
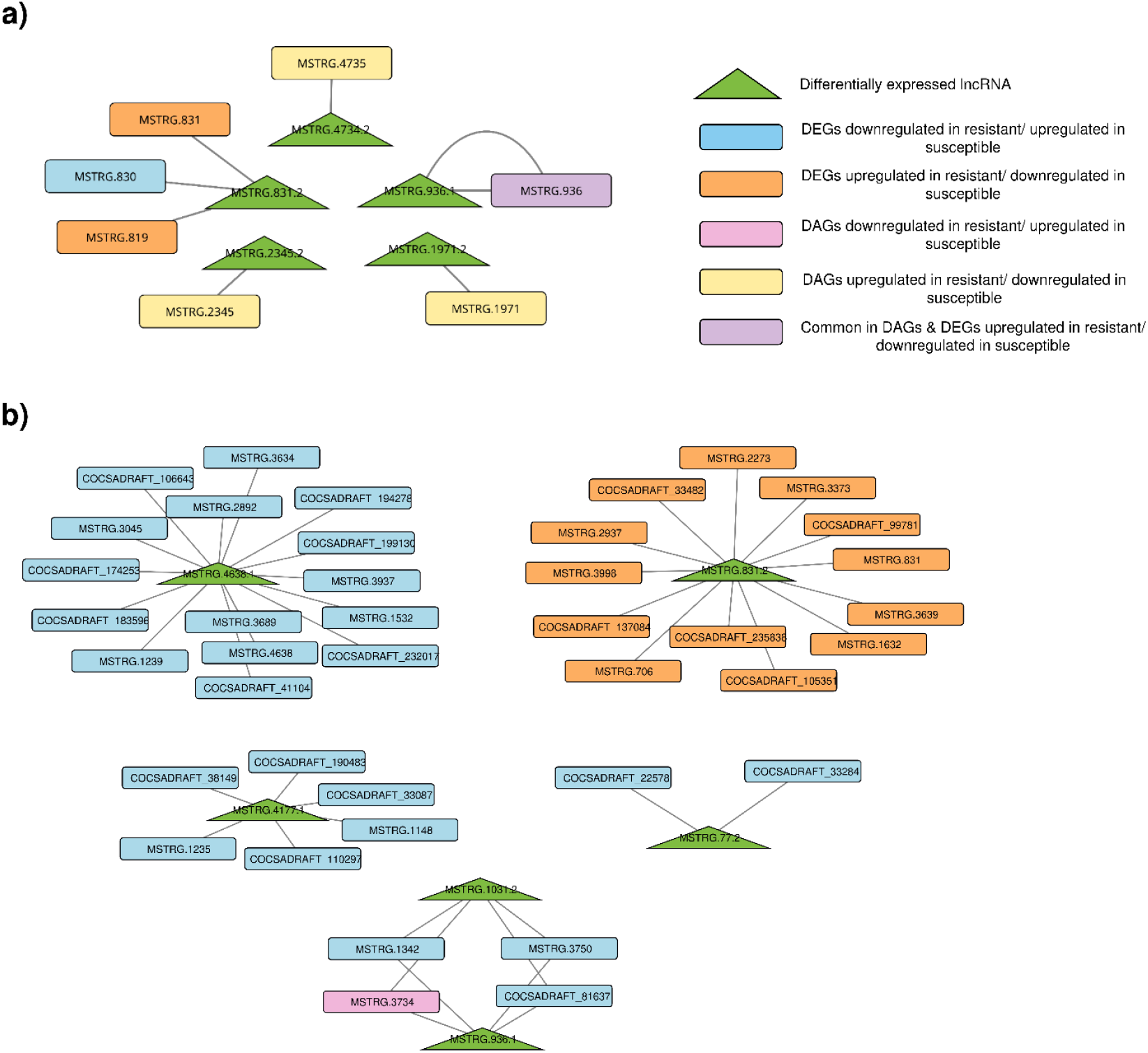
Integrated regulatory network of DEGs, DAS genes, and DElncRNAs in *Bipolaris sorokiniana*. The network illustrates potential cis- and trans-regulatory interactions among DEGs, differentially alternatively spliced (DAS) genes, and differentially expressed long non-coding RNAs (DElncRNAs) during infection of resistant and susceptible wheat genotypes. a) Cis-targets were identified based on genomic proximity (±100 kb), while b) trans-targets were predicted using Spearman correlation (r ≥ 0.9, p < 0.05). Visualization was performed in Cytoscape to uncover co-regulated modules and transcriptional coordination across regulatory layers.

GO enrichment analysis of cis-regulated target genes revealed strong enrichment in pathways related to RNA modification, RNA processing, and spliceosomal complexes, including the Box H/ACA snoRNP complex, U12-type spliceosome, and snRNA pseudouridine synthesis, suggesting a role for cis-acting lncRNAs in regulating RNA metabolism and post-transcriptional control (Fig. S6a). In contrast, trans-regulated targets were enriched in categories related to the Golgi apparatus, vesicle-mediated transport, and partially uncharacterized functions, including diacylglycerol binding and DNA replication regulation. These findings indicate that cis-acting lncRNAs may modulate core RNA processing machinery. In contrast, trans-acting lncRNAs influence broader cellular functions, including trafficking and cell cycle control, potentially contributing to host-specific fungal adaptation (Fig. S6b).

Among the trans-acting DElncRNAs, two fungal DELncRNAs, MSTRG.4638.1 and MSTRG.831.2, emerged as key regulatory elements. MSTRG.4638.1 exhibited exclusive interaction with fungal DEGs that were DownR/UpS, indicating a potential role in promoting pathogenicity or host susceptibility. GO enrichment analysis of its targets revealed significant enrichment for epigenetic and nuclear functions, such as histone methylation, DNA replication origin binding, chromatin binding, and protein alkylation. Additional enrichment in Golgi-associated vesicles and coated vesicles suggests a role in secretory and trafficking processes, potentially linked to effector delivery or host manipulation. In contrast, MSTRG.831.2 interacted predominantly with fungal genes that were UpR/DownS, implying a possible contribution to fungal adaptation or stress response during host defense. GO enrichment of its targets identified highly significant terms, including diacylglycerol binding and Golgi cis-cisterna organization, ATPase activator activity, and multivesicular body formation. Several categories, though currently uncharacterized, were suggestive of their roles in membrane dynamics, mitotic regulation, and auxin-responsive pathways, possibly reflecting adaptive strategies of the fungus under host-imposed stress.

## 4. Discussion

Understanding how fungal pathogens reprogram their transcriptional and post-transcriptional machinery during host colonization is critical to unravel the molecular basis of virulence (Rana et al. 2021; Mapuranga et al. 2023). *Bipolaris sorokiniana* is a globally significant hemibiotrophic pathogen, responsible for spot blotch, root rot, crown rot, and black point diseases in wheat and barley, leading to considerable yield and quality losses (Al-Sadi 2021). The pathogen’s broad host range and ability to infect multiple plant organs underscore its adaptive success and the complexity of its interaction. However, its infection-stage regulatory mechanisms are not yet well characterized (Al-Sadi 2021; Yuan et al. 2025). Recent high-quality genome assemblies of *B. sorokiniana* revealed an extensive repertoire of protein-coding genes, including hundreds of predicted secreted proteins and effectors, highlighting the molecular arsenal available to *B. sorokiniana* for host colonization and immune evasion (Yadav et al. 2023; Zhang et al. 2023). While advances in transcriptome profiling have deepened our understanding of host plant responses during infection, pathogen-centered studies remain sparse, particularly those examining *in planta* fungal responses. Ye et al. (2019) performed a comparative transcriptome and metabolomics analysis of *B. sorokiniana* infecting wheat to reveal host-induced transcriptional shifts, while Yazawa et al. (2013) and Mizuno et al. (2012) analyzed the simultaneous transcriptomes of *Bipolaris sorghicola* and sorghum to identify putative key factors in the interaction. A more recent study by Basak et al. (2024) employed dual RNA-seq to profile both *B. sorokiniana* and its host barley, identifying early-expressed fungal effectors and host defense transcripts. Similarly, Zhang et al. (2022) functionally characterized a novel effector, *CsSp1*, from *B. sorokiniana*, which is critical for wheat root rot. Additionally, Meshram et al. (2022) investigated *B. maydis* pathogenicity in maize. However, these studies primarily focused on effector identification or early host responses without exploring pathogen-centered regulatory layers, such as AS and lncRNAs, particularly within the context of infection *in planta*.

In this context, our study provides a comprehensive *in planta* gene regulation overview of *B. sorokiniana* by integrating differential gene expression, alternative splicing, and lncRNA analyses during infection of spot blotch-resistant and susceptible wheat genotypes. This multifaceted approach captured the pathogen’s active transcriptional programs and highlighted how RNA-level plasticity may facilitate its adaptation to host-imposed challenges. By focusing on the response of *B. sorokiniana* to contrasting immune environments, our study offers novel insights into the pathogen’s molecular strategies to promote infection and overcome host resistance barriers. The integrated transcriptome analysis revealed the pathogen’s host-genotype-dependent transcriptional reprogramming, distinguishing successful colonization in the susceptible wheat genotype from the suppressed pathogenicity in the resistant wheat genotype.

### 4.1 Host-dependent transcriptional shifts in *B. sorokiniana*

Transcriptomic profiling of *B. sorokiniana* during infection of resistant and susceptible wheat genotypes revealed pronounced host genotype-dependent gene expression dynamics, underscoring the pathogen’s transcriptional adaptability to distinct immune environments. A total of 128 fungal genes were significantly upregulated in the susceptible host (DownR/UpS), leading to aggressive fungal colonization. Functional enrichment analyses revealed a marked overrepresentation of ribosome biogenesis, spliceosome-associated components, RNA splicing factors, and mRNA processing regulators. This suggests a coordinated upregulation of post-transcriptional and translational machinery, reflecting greater biosynthetic activity that might have enhanced fungal proliferation and virulence in the susceptible host. Such transcriptional signatures are hallmarks of rapid growth and biomass accumulation (Woolford and Baserga 2013).

Ribosome biogenesis plays a central role in meeting the protein synthesis demands of infection (Bhabhra et al. 2008). Specifically, the upregulation of *ribosome biogenesis AAA ATPase RIX7* (MSTRG.328) and *GAR1* (MSTRG.936) domain-containing protein in the susceptible genotype likely reflected a metabolic environment conducive to fungal proliferation. RIX7 is part of the essential AAA ATPase family involved in the maturation of the 60S ribosomal subunit, a process critical for assembling functional ribosomes. In *Saccharomyces cerevisiae*, RIX7 was shown to be indispensable for ribosomal export from the nucleus (Gadal et al. 2001). Its homolog in *C. albicans*, CaYLL34, complements defective *rix7* mutants, indicating conserved functionality (Puzer et al. 2006). Beyond its role in ribosome assembly, *CaYLL34p* was implicated in morphogenetic switching, linking ribosome biogenesis to pathogenic development in fungi (Puzer et al. 2006). These findings indicate that RIX7-like ATPases sustain translational capacity and interface with broader virulence-related processes.

Similarly, GAR1 is a core component of the H/ACA small nucleolar ribonucleoprotein (snoRNP) complex, which mediates pseudouridylation of rRNA; a crucial RNA modification required for ribosome stability and function (Watkins and Bohnsack 2012; Czekay and Kothe 2021). In fungal systems, the accurate folding and maturation of rRNAs under normal and stress conditions are essential to maintain protein synthesis during infection. Studies in yeast have shown that mutations in components of the H/ACA snoRNP complex, such as SHQ1, impair snoRNA expression and rRNA processing, resulting in growth defects. This highlights the essential role of H/ACA snoRNPs in ribosome biogenesis and cellular fitness (Grozdanov et al. 2009). Interestingly, it has been demonstrated that disruption or silencing of H/ACA-class snoRNAs interferes with viral replication, as seen in Epstein–Barr virus and respiratory viruses like SARS-CoV and influenza, suggesting broader roles of these snoRNAs in stress responses and pathogenic regulation (Murray et al. 2014). In our study, the upregulation of fungal *GAR1* in the susceptible host suggests an enhanced translational machinery that enables the fungus to exploit weakened host defenses. Given the pivotal role of ribosome biogenesis in cellular proliferation, its misregulation can impair fungal growth and survival (Bhabhra et al. 2008; Woolford and Baserga 2013). These findings highlight how *B. sorokiniana* reprograms core cellular pathways to favor pathogenic success, and position RIX7 and GAR1 as candidate molecular hubs with potential for antifungal intervention.

Functional genomics analysis of *B. sorokiniana* during infection of resistant (RG) and susceptible (SG) wheat genotypes revealed a striking dichotomy in the pathogen’s adaptive strategies. GO enrichment analysis showed that DownR/UpS genes were predominantly associated with ribosome biogenesis, spliceosomal function, carbohydrate metabolism, and oxidative stress response, suggesting that the pathogen actively maintained high levels of protein synthesis and RNA processing to facilitate rapid growth and colonization in the susceptible host. Conversely, UpR/DownS genes were enriched in GO categories associated with cell wall biosynthesis, intracellular transport, and lipid metabolism, including sphingolipid biosynthesis and vesicle-mediated trafficking. These findings suggest that the resistant host environment imposed a metabolic bottleneck, reducing the pathogen’s protein synthesis capacity and cellular proliferation, and forcing it to strengthen its cellular integrity and sustain survival.

Further, protein-protein interaction (PPI) network revealed host-genotype-specific functional clustering, offering deeper mechanistic insights into how the pathogen orchestrates infection strategies under varying immune pressures. The susceptible host background showed a densely saturated PPI network enriched in proteins associated with the spliceosomal complex, mRNA processing, and RNA splicing. This suggests a robust transcriptional and post-transcriptional regulatory program facilitating efficient RNA maturation and protein synthesis during infection. Such enrichment is consistent with a hyperactivated biosynthetic state required for aggressive colonization and fungal proliferation. Though smaller, the UpR/DownS gene set revealed enrichment of sphingolipid biosynthesis and sulfate assimilation pathways, reflecting an adaptive fungal response to the resistant host environment. These results highlight a clear functional divergence in *B. sorokiniana*’s transcriptional response driven by the host environment. Infection of the susceptible host is characterized by increased RNA processing and post-transcriptional activity, potentially supporting rapid fungal growth and effector regulation. Conversely, upregulation of lipid metabolism-related genes in the resistant host may represent a fungal counter-defense strategy to maintain cellular integrity and mitigate host-imposed stress.

Interestingly, this molecular process mirrors the mechanisms observed in other phytopathogenic fungi. For example, in *F. graminearum*, the spliceosome-associated kinase *FgPrp4* plays a critical role in B-complex activation and overall splicing efficiency, with its loss resulting in impaired development and pathogenicity (Gao et al. 2016). Similarly, in rust fungi, effector proteins directly bind to host pre-mRNA splice sites, thereby reprogramming AS to suppress host resistance (Tang et al. 2022). Collectively, these findings underscore a conserved strategy across fungal pathogens, centered on RNA processing and membrane maintenance to overcome host resistance.

The enrichment of sphingolipid and ceramide metabolism pathways is particularly interesting, as these play roles in fungal stress signaling and programmed cell death, enabling the pathogen to tolerate host-induced stress better (Munshi et al. 2018; Mota Fernandes and Del Poeta 2020; Wang et al. 2021; Li et al. 2022b). The enrichment of lipid metabolic and cell wall biosynthesis pathways in resistant interactions suggests that *B. sorokiniana* actively responds by reinforcing its cellular structures and altering membrane composition. The CAZy annotations revealed that 34 of the 42 differentially expressed carbohydrate-active enzymes (CAZymes) were upregulated in the susceptible host. These predominantly included glycoside hydrolases, glycosyltransferases, and auxiliary activity enzymes, which degrade plant cell walls, facilitating host penetration and nutrient assimilation (Zhang et al. 2020; Li et al. 2022c). Their co-enrichment within the susceptible host indicates a coordinated enzymatic attack on host structural barriers, synchronized with high transcriptional activity. *Bipolaris sorokiniana* utilizes a variety of CAZymes to degrade host cell walls and obtain nutrients. This behavior aligns with virulence strategies commonly observed in necrotrophic and hemibiotrophic fungi (Lyu et al. 2015; Ramzi et al. 2019). Consistent with this, PHI-base uncovered 34 DEGs with homology to known virulence factors, of which 20 were upregulated in the susceptible background. This supports the hypothesis that these genes are central to the pathogen’s virulence strategy. *B. sorokiniana* may actively utilize these virulence determinants during infection of susceptible wheat to suppress immune responses or manipulate host metabolism (Aoun 2017). In contrast, genes associated with reduced virulence or unaffected pathogenicity were upregulated in the resistant background, hinting at a suppression or misregulation of core pathogenicity components under resistance-imposed stress. This underscores the transcriptional flexibility of the pathogen in deploying or downregulating virulence strategies based on the host genotype. Exclusive upregulation of predicted effectors in the susceptible host further underlines the pathogen’s deployment of host-manipulating molecules to facilitate colonization and evade immune detection.

### 4.2 Alternative splicing as a regulatory level in *Bipolaris sorokiniana* pathogenicity

Alternative splicing (AS) is increasingly recognized as a key mechanism by which eukaryotic pathogens diversify their transcriptomes, allowing them to rapidly and precisely modulate gene expression in response to environmental stimuli. Traditionally thought to be infrequent in fungi, emerging evidence has now established that AS is widespread and functionally significant in several fungal species, including both human and plant pathogens (Grützmann et al. 2014; Gonzalez-Hilarion et al. 2016; Muzafar et al. 2021). These AS events play pivotal roles in fungal growth, development, stress adaptation, and potentially in pathogenicity and drug resistance.

Our study provides the first *in planta* evidence of AS in *B. sorokiniana* during its interaction with wheat. A total of 116 AS events were detected, with 22 differentially regulated between infections of resistant (RG) and susceptible (SG) wheat genotypes. Intron retention (RI) and alternative donor or acceptor splice site usage were the most dominant AS types. Thus, the pathogen probably uses these events to fine-tune gene expression in response to contrasting host environments. These patterns are consistent with reports in *C. neoformans* and *S. sclerotiorum*, where retained introns and altered splice sites were also the most prevalent AS forms (Gonzalez-Hilarion et al. 2016; Ibrahim et al. 2021). The differential regulation of AS events between resistant and susceptible wheat hosts detected in this study, points toward a host-genotype-dependent control of splicing in *B. sorokiniana*, enabling the fungus to remodel its secretome or stress-response repertoire under varying immune environments. AS rates in fungal pathogens tend to be modest compared to higher eukaryotes, with estimates ranging from 0.6% in *Colletotrichum graminicola* (Schliebner et al. 2014) to 38.82% in *Histoplasma capsulatum* (Muzafar et al. 2021). The AS frequency observed in *B. sorokiniana* aligns with this range, 0.95%, suggesting a regulatory rewiring of the fungal transcriptome in response to environmental cues. Such modulation may enhance virulence in permissive environments while enabling stress adaptation under hostile conditions.

Functionally, AS events such as alternative 5’ splice sites (A5SS) and alternative last exons (ALE) were predominantly associated with genes involved in translation, suggesting a role in modulating protein synthesis to suit the host environment. Notably, the differentially alternatively spliced gene *MSTRG*.*4962*, which encodes a PABP domain-containing protein, exhibited an A5SS event and was upregulated in the susceptible host but downregulated in the resistant genotype. Annotation against the PHI-base revealed its orthologs in *C. neoformans, U. maydis*, and *F. graminearum* to be linked to reduced virulence or loss of pathogenicity phenotypes. This gene is implicated in mRNA metabolism, stability, and translational efficiency. In *S. cerevisiae*, disruption of its homolog severely impaired cell growth, underscoring its essential role in post-transcriptional regulation and fungal fitness (Sachs et al. 1987). A3SS (Alternative 3⍰ Splice Site) and AFE (Alternative First Exon) targeted transcripts linked to endoplasmic reticulum function, macromolecular metabolism, and secondary biosynthesis pathways critical for stress response and virulence modulation (Zhao et al. 2013; Grützmann et al. 2014; Jin et al. 2017).

Intron retention (IR) is the most common form of alternative splicing observed in fungi and is frequently associated with transcriptional regulation under stress conditions (Fang et al. 2020). In *B. sorokiniana*, differential IR events were identified in key genes, including *MSTRG*.*1971* (*methionine aminopeptidase*) and *MSTRG*.*4735* (*plasma membrane H*^*+*^*-ATPase*), both of which are crucial for fungal growth, stress adaptation, and pathogenicity. Methionine aminopeptidases (MetAP), beyond their conserved role in N-terminal protein processing, have been linked to virulence across kingdoms, for instance, as secreted effectors in *Citrobacter rodentium* that target host mitochondria (Xia et al. 2019). Inhibition of MetAP in bacteria has been shown to halt growth (Helgren et al. 2016). In fungal pathogens, it is critical for the post-translational processing of proteins, supporting fungal growth, survival, and likely virulence. Likewise, fungal H^+^-ATPases are key regulators of nutrient uptake and pH homeostasis. Their disruption resulted in a complete loss of pathogenicity in *Leptosphaeria maculans* (Remy et al. 2008a) and *Colletotrichum higginsianum* (Korn et al. 2015) and led to reduced virulence in *F. graminearum* (Wu et al. 2022). IR in such genes may have a regulatory function, enabling pathogens to fine-tune growth and host adaptation during infection. Another intron retention differentially alternative spliced gene *MSTRG*.*385*, encodes a serine/threonine-protein phosphatase, indicative of post-transcriptional regulation that may modulate its expression under specific host conditions.

Serine/threonine phosphatases are evolutionarily conserved enzymes that catalyze the removal of phosphate groups from phosphoserine/threonine residues; thereby regulating a wide range of cellular processes, including cell cycle progression, stress signaling, morphogenesis, and virulence (Ariño et al. 2019). In *M. oryzae*, deletion of the PP2A catalytic subunit MoPPG1 severely impairs vegetative growth, conidiation, appressorium formation, and abolishes pathogenicity, highlighting its central role in infection-related development (Osés-Ruiz et al. 2021). Similarly, in *Fusarium oxysporum*, the PP2C-type phosphatase Ptc6 is critical for MAPK pathway regulation and is required for host penetration and colonization, underlining its involvement in stress adaptation and virulence (Nunez-Rodriguez et al. 2020). Consistently, PHI-base annotation of MSTRG.385 aligns it with orthologs in *C. albicans, F. graminearum*, and *M. oryzae*, all associated with reduced virulence phenotypes. These findings suggest that alternative splicing of this phosphatase gene may fine-tune its activity as part of a broader fungal adaptation strategy to host resistance pressure.

Notably, two differentially alternatively spliced genes were associated with the spliceosomal machinery itself. *MSTRG*.*2345*, encoding small nuclear ribonucleoprotein SmD3, was upregulated in the resistant genotype, whereas *MSTRG*.*936*, encoding GAR1, a core component of the H/ACA snoRNP complex, was significantly upregulated in the susceptible genotype. SmD3 is a core component of the U1, U2, U4/U6, and U5 snRNPs, forming the essential Sm protein ring crucial for spliceosome assembly and function (Will and Lührmann 2011). Perturbation in SmD3 expression can destabilize snRNP formation and reduce splicing efficiency. For example, in *Arabidopsis thaliana*, disruption of SmD3 compromised both constitutive and alternative splicing mechanisms. It enhanced susceptibility to *Pseudomonas syringae*, due to impaired stomatal immunity and altered expression of defense-related transcripts. Upregulation of SmD3 in *B. sorokiniana* during resistant host interaction might reflect an adaptive mechanism to maintain splicing fidelity under resistant host pressure, or it could indicate a broader stress response regulatory role during infection. Its involvement in post-transcriptional control could influence fungal gene expression reprogramming required for survival under host-induced stress. Likewise, GAR1, part of the H/ACA snoRNP complex, is pivotal to pseudouridylation of rRNA, a modification critical for ribosome maturation and fidelity (Watkins and Bohnsack 2012; Czekay and Kothe 2021). Its dual categorization as a DAG and DEG reinforces its centrality in coordinating transcriptional, post-transcriptional, and translational layers of regulation. The upregulation of GAR1 in the susceptible host interaction may enhance ribosomal efficiency to meet the elevated translational demands of infection.

### 4.3 Emergence of lncRNAs as regulatory elements in fungal pathogenicity

lncRNAs are ubiquitous across all domains of life. Although extensively characterized in mammals, they are also prevalent in plants and fungi, including the model yeasts *S. cerevisiae* and *Schizosaccharomyces pombe* (Chacko and Lin 2013; Li et al. 2021; Mattick et al. 2023). The latest research has revealed their pivotal functions in pathogens, particularly in phytopathogenic fungi. The genus *Fusarium*, for example, has been a rich source for lncRNA characterization. Studies in *F. graminearum* demonstrated that lncRNA expression is linked to fungal development and infection-related morphogenesis, with lncRNAs like lncRsp1 and lncRsn directly contributing to virulence on wheat (Kim et al. 2018; Wang et al. 2022; Fu et al. 2024). However, this regulatory role is not limited to virulence, as seen with the carP lncRNA modulating carotenoid biosynthesis in *F. oxysporum* and *F. fujikuroi* (Parra-Rivero et al. 2020). While these studies establish lncRNAs as key regulators, their roles in the *B. sorokiniana*-wheat pathosystem remain uncharacterized.

Beyond protein-coding genes and canonical splicing components, lncRNAs are increasingly recognized as key regulators of fungal gene expression and virulence. In particular, they may interface with transcriptional and splicing processes to coordinate dynamic gene regulation in response to host-derived cues. Higher expression of lncRNAs during infection of the susceptible host genotype suggests that they could function as pathogenic factors, with their expression suppressed during a successful host defense response but induced to promote infection in a susceptible host. This host-dependent regulation of fungal lncRNAs aligns with previous findings in other plant pathogens. For instance, dynamic lncRNA expression has been reported in *V. dahliae* during cotton infection (Li et al. 2022a) and has been associated with the development of infection structures in *M. oryzae* and *Botrytis cinerea* (Choi et al. 2022; Shi et al. 2025). Furthermore, in *Zymoseptoria tritici*, lncRNAs are predominantly expressed during early infection and are often co-expressed with stress-responsive and pathogenicity-related genes (Glad et al. 2023), supporting their role in fungal adaptation and virulence. In contrast to this general trend, MSTRG.831.2 was the only lncRNA that was upregulated in the resistant interaction (UpR/DownS) in this study. Target prediction analysis indicated that MSTRG.831.2 may regulate trans target *COCSADRAFT_33482*, a gene encoding a Mon1 homolog. In *S. cerevisiae, Mon1* is a key protein involved in cytoplasm-to-vacuole trafficking and vacuolar fusion, processes essential for cellular homeostasis and stress survival (Jensen 2016). This function is conserved in pathogenic fungi. Deletion of *Mon1* homologs in *M. oryzae* and *F. graminearum* impairs vacuolar assembly, autophagy, and pathogenesis (Gao et al. 2013; Li et al. 2015). The specific upregulation of MSTRG.831.2 in the resistant host genotype suggests it may function as part of a host-induced mechanism that disrupts *Mon1*-dependent pathways, thereby impairing fungal viability and contributing to disease resistance.

Among the DElncRNA, MSTRG.4638.1 was identified as a potential regulatory hub, predicted to control multiple trans-targets with established roles in fungal virulence. Its key trans-target is COCSADRAFT_41104, which encodes a Glycoside Hydrolase family 61 (GH61) protein, also known as a lytic polysaccharide monooxygenase (LPMO). LPMOs are critical for degrading plant cell wall components, facilitating fungal penetration (Westereng et al. 2011; Rafiei et al. 2021). The strong downregulation of this gene in the resistant genotype (log_2_FC = - 5.12) and its annotation as a predicted apoplastic effector underscore its likely importance in host colonization. MSTRG.4638.1 is also predicted to regulate genes essential for fundamental cellular processes. These include MSTRG.3689, which encodes the pre-mRNA-splicing factor Clf1, whose deletion in *C. neoformans* reduced virulence (Chung et al. 2003), and MSTRG.3045, encoding the cytoskeletal protein α-tubulin (TUB1), which is essential for pathogenicity in *Aspergillus fumigatus* (Hu et al. 2007). The coordinated upregulation of MSTRG.4638.1 and these trans-targets in the susceptible host suggests this lncRNA facilitates infection by enhancing the efficiency of gene processing and cytoskeletal dynamics required for invasive growth. Furthermore, MSTRG.4638.1 targets COCSADRAFT_106643, a FYVE zinc finger-domain-containing protein. Homologs like PsFP1 in *Phytophthora sojae* are required for virulence and oxidative stress tolerance (Zhang et al. 2021; Ru et al. 2023). This regulatory axis may help the pathogen cope with host-induced stress. In another regulatory mechanism, MSTRG.4734.2 acts in *cis* on its neighboring gene, MSTRG.4735, encoding a plasma membrane ATPase. The homolog of this gene in *Leptosphaeria maculans* is essential for pathogenicity, highlighting the importance of maintaining ion homeostasis and membrane potential during host invasion (Remy et al. 2008b). The potential regulation of this ATPase by MSTRG.4734.2 suggests a direct role of the lncRNA in modulating fungal membrane physiology and virulence.

In summary, this study demonstrates that lncRNAs in *B. sorokiniana* could function as key regulatory molecules that respond dynamically to the host environment. We identified a predominant class of lncRNAs, exemplified by MSTRG.4638.1, that appear to function as potential pathogenic factors by coordinating the expression of multiple virulence-associated genes. Conversely, the resistant-specific expression of lncRNA MSTRG.831.2, coupled with its predicted trans-interactions with multiple target genes, including Hsp90 ATPase activators, cytochrome C oxidase assembly protein (COX15), and trafficking regulators like Mon1, suggests a potential role in modulating mitochondrial function, protein homeostasis, and vesicle-mediated signaling under host-imposed stress. Notably, its interaction with a dDENN domain-containing protein (COCSADRAFT_99781), previously found upregulated in the resistant interaction, points toward a role in regulating vesicular trafficking and membrane dynamics during defense. These interactions position MSTRG.831.2 as a putative regulatory hub in fungal adaptation to resistant host environments.

### 4.4 Spliceosomal modulation as a fungal adaptation strategy

To visualize the pathogen’s post-transcriptional regulatory shifts, we identified differentially expressed spliceosome-related genes of the canonical spliceosome pathway (Fig. 6a). This revealed coordinated modulation of five core components, PRP8BP, SF3A, AQR, Syf, and SmD3 in *B. sorokiniana* during infection of susceptible wheat genotype, suggesting an adaptive restructuring of the splicing machinery in response to the host environment. PRP8BP (Pre-mRNA Processing Factor 8 binding protein) is a central scaffold protein of the spliceosome, essential for both its assembly and catalytic function. Disruption of PRP8 impairs pre-mRNA processing and broadly impacts gene expression (Grainger and Beggs 2005). Notably, in fungal pathogens like *C. neoformans*, PRP8 contains a self-splicing intein whose inhibition by metal ions or antifungal agents can suppress fungal growth, highlighting its therapeutic potential (Green et al. 2019; Li et al. 2021). SF3A, a subunit of the U2 snRNP complex, plays a critical role in intron recognition and has been linked to morphogenesis and virulence in *C. albicans*, where its disruption hinders the yeast-to-hyphal transition essential for infection (Lash et al. 2024). Structural studies also underscore its conserved function across fungi in maintaining spliceosome integrity (Lin and Xu 2012).

**Fig. 6.**
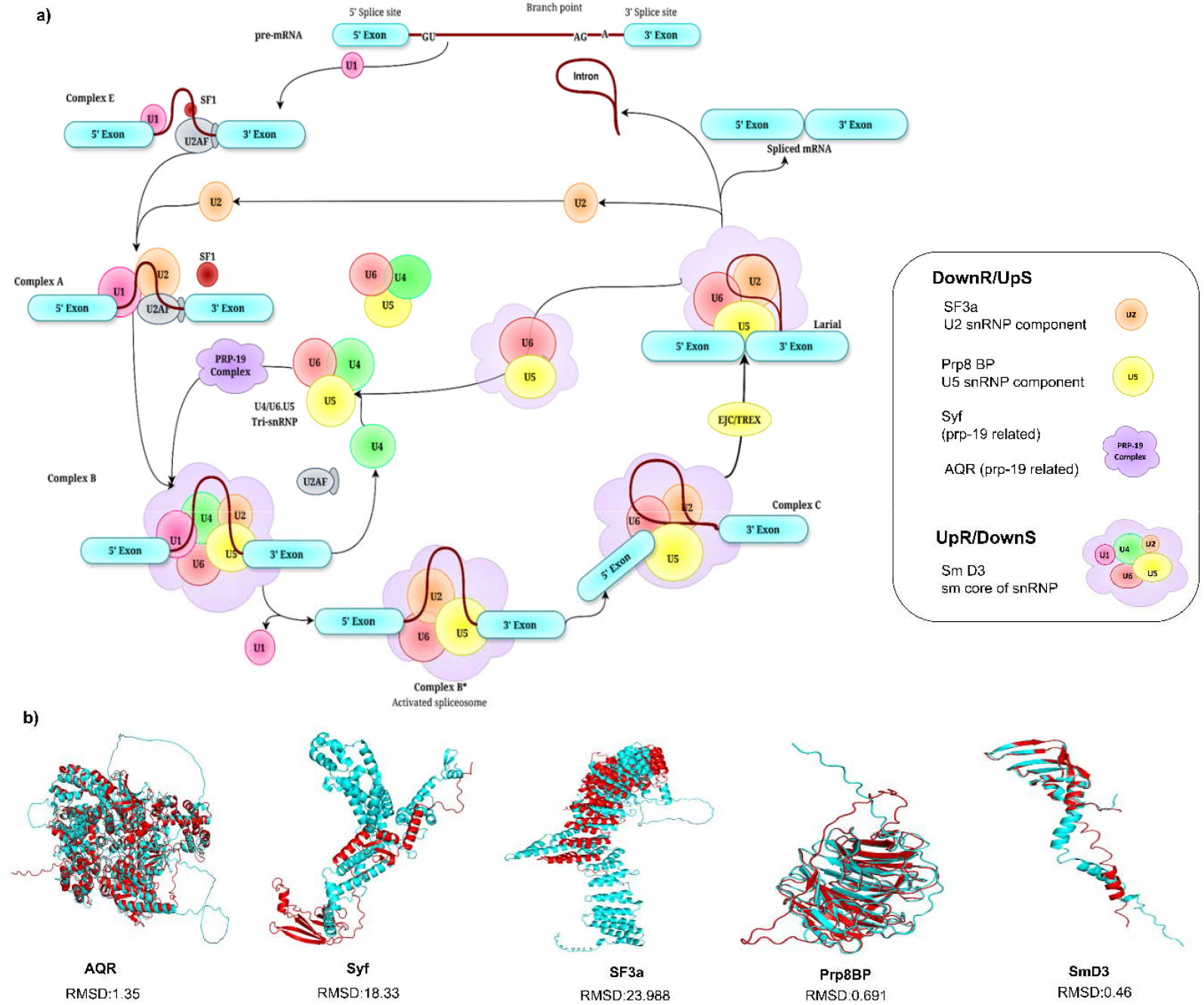
Spliceosomal modulation and structural divergence in *Bipolaris sorokiniana* during wheat infection. (a) Canonical spliceosome pathway of *B. sorokiniana* depicting the differential regulation of five core spliceosomal components—PRP8BP, SF3A, AQR, Syf, and SmD3—during infection of resistant and susceptible wheat genotypes. These genes show coordinated modulation, suggesting a host-dependent remodeling of the fungal splicing machinery. (b) Structural superimposition of five key spliceosomal proteins from *B. sorokiniana* and their wheat homologs reveals marked conformational divergence, particularly in SF3A and Syf. Structures were predicted using SWISS-MODEL and visualized in PyMOL, highlighting potential differences in protein-protein or protein-RNA interaction interfaces that may be exploited for pathogen-specific antifungal targeting.

AQR (Aquarius) is an intron-binding spliceosomal factor involved in maintaining splicing fidelity by preventing cryptic splice site usage (Li 2020). Its downregulation during the resistant host interactions may reflect a tighter control on transcriptome remodeling under resistance pressure. Syf, a suppressor-of-forked domain protein, contributes to spliceosome stability and is implicated in co-transcriptional splicing in yeast (Will and Lührmann 2011), although its role in filamentous fungi remains underexplored. SmD3, a conserved spliceosomal core component, emerged as a potential virulence-associated factor in *B. sorokiniana*, exhibiting AS and significant upregulation in the resistant wheat genotype, indicating host-specific regulation. Since SmD3 is essential for spliceosome assembly and pre-mRNA splicing, its differential regulation suggests a dynamic role in post-transcriptional adaptation during host-pathogen interaction. Although direct functional characterization of SmD3 in fungal pathogens is limited, its conserved role in splicing across eukaryotes, and the *Arabidopsis* study showing that SmD3 disruption compromises resistance and alters alternative splicing of resistance-related transcripts (Golisz et al. 2021) underscores its relevance.

The differential regulation of these spliceosomal components reflects a strategic reprogramming of *B. sorokiniana*’s RNA processing machinery to optimize gene expression in response to host susceptibility. These findings suggest that spliceosome dynamics not only support general RNA metabolism but may also be leveraged by the pathogen to express host-adaptive and virulence-associated transcripts, making them promising targets for future antifungal interventions.

### 4.5 Structural divergence of fungal spliceosomal proteins highlights potential for pathogen-specific targeting

To evaluate the therapeutic potential of targeting spliceosomal components in *B. sorokiniana*, we performed structural comparisons of key spliceosome-related proteins between the pathogen and its host, wheat, using PyMOL-based 3D superposition (Fig. 6b). Homology modeling with SWISS-MODEL (Waterhouse et al. 2018) revealed that while sequence identity ranged from 18% to 42% for DEGs and up to 60% for differentially alternatively spliced genes, the root mean square deviation (RMSD) values, particularly for SF3A and SYF proteins, were substantially high, indicating marked structural divergence. Despite conservation of certain core elements, the superimposed models showed significant conformational shifts in key domains, suggesting potential alterations in protein-protein or protein-RNA interactions. Given the complexity and dynamic regulation of the spliceosome, even minor architectural variations can lead to altered splicing activity and substrate specificity (Will and Lührmann 2011; Papasaikas and Valcárcel 2016). This structural flexibility may underlie the ability of *B. sorokiniana* to fine-tune its transcriptome in a host-dependent manner, aiding its pathogenic adaptability. Importantly, the fungal-specific divergence in these essential proteins opens a therapeutic window for selective inhibition, minimizing off-target effects in the host. This concept is supported by prior antifungal strategies targeting the Prp8 intein in *C. neoformans* (Li et al. 2022a). Thus, our findings suggest that the evolutionary divergence of spliceosomal proteins in *B. sorokiniana* not only facilitates niche-specific transcriptomic remodeling but also offers a promising molecular basis for pathogen-specific intervention strategies.

## 5. Conclusions

This study provides the first comprehensive *in planta* dissection of *Bipolaris sorokiniana*’s transcriptional and post-transcriptional responses during infection of contrasting wheat genotypes. This study reveals a dynamic, host-genotype-dependent reprogramming of the fungal transcriptome by integrating various analyses, such as dual RNA-seq, differential gene expression, alternative splicing, and identification of lncRNAs. In the susceptible wheat variety, *B. sorokiniana* exhibited enhanced expression of genes related to ribosome biogenesis, RNA processing, and metabolic activity, reflecting a transcriptional architecture optimized for aggressive colonization. In contrast, resistant host interaction was characterized by suppression of the core biosynthetic functions of the fungal pathogen and enriching its stress-adaptive responses. Spliceosomal pathway genes, such as *PRP8BP, SF3A, AQR, Syf, SmD3*, and ribosome biogenesis genes like *AAA ATPase RIX7* and *GAR1*, along with lncRNAs, MSTRG.4638.1 and MSTRG.831.2, emerged as central regulators of transcriptomic adaptation, influencing splicing, protein translation, and virulence-associated pathways. Furthermore, structural modeling revealed significant divergence between fungal and wheat spliceosomal proteins, especially SF3A and SYF, suggesting their potential as selective antifungal targets. Our findings demonstrate that *B. sorokiniana* leverages transcriptional flexibility, AS, and lncRNA-mediated regulation to adapt to immune environments and modulate virulence. These insights provide a foundation for targeted functional studies and open new avenues for host-specific antifungal strategies to disable key post-transcriptional regulators without affecting the host.

## Acknowledgements

**BRD** acknowledges the University Grants Commission (UGC), India, for the junior and senior research fellowships and the Academy of Scientific and Innovative Research (AcSIR), India, for enrollment in the Ph.D. program. **AKS** acknowledges AcSIR, India, for the Integrated Dual-Degree Program (IDDP) and the research fellowship for IDDP under the CSIR-GATE-JRF program (Award No.: 31/ GATE/11(55)/2025-EMR-I). **KRK** is thankful for the research fellowship and grants from the Department of Science and Technology (DST)-INSPIRE Faculty Program (Award number: IFA18-LSPA123). The authors acknowledge Ms. Ashwini Gudekar, CSIR-NCL, Pune, for her contribution as a project intern and for generating Fig. 3a and 6a used in this study. This research was funded by the Council of Scientific and Industrial Research (CSIR), India (Grant number: MLP101226).

## Author Contributions

**Bhakti R. Dayama**: Conceptualization, Data curation, Formal analysis, Investigation, Methodology, Visualization, Writing-original draft. **Anand Kumar Shukla**: Conceptualization, Investigation, Methodology, Formal analysis, Visualization, Writing-original draft. **Kirtikumar R. Kondhare**: Conceptualization, Resources, Data curation, Supervision, Writing-review and editing. **Narendra Y. Kadoo**: Conceptualization, Funding acquisition, Project administration, Resources, Supervision, Writing-review and editing.

## Data availability statement

The authors declare that the data supporting the findings of this study are available within the manuscript. The RNA-sequencing raw reads have been deposited in the Sequence Read Archive (SRA) of the National Center for Biotechnology Information (NCBI) database (https://www.ncbi.nlm.nih.gov/sra) with the accession number PRJNA1090403.

## Supplementary Figure Legend

**Fig. S1. Gene Ontology annotations of DEGs of *Bipolaris sorokiniana* during infection of resistant and susceptible wheat genotypes**. The bar chart presents enriched Gene Ontology (GO) terms under biological process, molecular function, and cellular component categories for genes upregulated in resistant (UpR/DownS) and susceptible (DownR/UpS) host backgrounds.

**Fig. S2. Functional categorization of *B. sorokiniana* DEGs reveals distinct GO categories during infection of resistant and susceptible wheat hosts**. Enriched GO terms for a) genes downregulated in resistant and upregulated in susceptible wheat genotypes (DownR/UpS), and b) genes upregulated in resistant and downregulated in susceptible wheat genotypes (UpR/DownS) were shown. The left panel displays fold enrichment (x-axis), where dot size reflects the number of DEGs associated with each GO term, and the dot color indicates – log10(FDR), the negative log-transformed false discovery rate from Fisher’s exact test. Higher log10(FDR) values denote stronger statistical significance. The right panels show hierarchical clustering of enriched GO terms, highlighting functional coherence and divergence of fungal responses in different host environments.

**Fig. S3. Functional features of *Bipolaris sorokiniana* differentially expressed genes**. Insights from (a) KEGG pathway analysis, (b) PHI-base database annotation, (c) CAZymes (Carbohydrate-Active enzymes) distributions, and (d) predicted effector proteins.

**Fig. S4. Gene Ontology enrichment of differentially alternatively spliced genes in *Bipolaris sorokiniana* during infection of resistant and susceptible wheat genotypes**. (a) differentially alternatively spliced genes (DAGs) downregulated in the resistant and upregulated in the susceptible wheat genotype (DownR/UpS). (b) DAGs upregulated in the resistant and downregulated in the susceptible wheat genotype (UpR/DownS).

**Fig. S5. Differential expression and characterization of long non-coding RNAs in *Bipolaris sorokiniana* during infection of resistant and susceptible wheat genotypes**. (a–b) Volcano plots showing differentially expressed long non-coding RNAs (lncRNAs) identified by DESeq2 and edgeR in resistant (RG) vs. susceptible (SG) wheat genotype. Significant lncRNAs (p-value < 0.2 and |log_2_FC| ± 2) are highlighted in red. (c) Venn diagram illustrating the overlap of differentially expressed lncRNAs identified by DESeq2 and edgeR. A total of 14 DElncRNAs were detected, with 13 showing consistent patterns of downregulation in RG and upregulation in SG. (d) Subcellular localization prediction of DElncRNAs, indicating their predominant localization to the cytoplasm, ribosome, and exosome, suggests potential regulatory roles at the post-transcriptional and translational levels. (e) Classification of DElncRNAs based on genomic context, showing the majority are sense lncRNAs, followed by antisense and intergenic types of lncRNAs.

**Fig. S6. Gene Ontology enrichment of predicted target genes regulated by differentially expressed long non-coding RNAs in *Bipolaris sorokiniana***. (a) Gene Ontology (GO) enrichment of cis-regulated target genes, and (b) GO enrichment of trans-regulated target genes.

## Supplementary Table Captions

**Table S1**: Summary of raw and quality-filtered RNA-seq data from *Bipolaris sorokiniana*-infected wheat leaf samples at 4 dpi

**Table S2**: Mapping and classification summary of *Bipolaris sorokiniana* reads extracted from dual RNA-seq data of infected wheat leaves at 4 dpi

**Table S3**: DESeq2: 162 differentially expressed genes from SI (Susceptible inoculated) vs RI (Resistant inoculated) comparison

**Table S4**: DESeq2: 116 genes downregulated in resistant and upregulated in susceptible (DownR/UpS)

**Table S5**: DESeq2: 46 genes upregulated in resistant and downregulated in susceptible (UpR/DownS)

**Table S6**: edgeR: 99 differentially expressed genes from SI (Susceptible inoculated) vs RI (Resistant inoculated) comparison

**Table S7**: edgeR: 64 genes downregulated in resistant and upregulated in susceptible (DownR/UpS)

**Table S8**: edgeR: 35 genes upregulated in resistant and downregulated in susceptible (UpR/DownS)

**Table S9**: Functional annotation of DEGs downregulated in resistant and upregulated in susceptible (DownR/UPS), including descriptions, GO terms, enzyme codes, and InterPro domains

**Table S10**: Functional annotation of DEGs upregulated in resistant and downregulated in susceptible (UpR/DownS), including descriptions, GO terms, enzyme codes, and InterPro domains

**Table S11**: KEGG pathways of DEGs downregulated in resistant and upregulated in susceptible (DownR/UpS)

**Table S12**: KEGG pathways of DEGs upregulated in resistant and downregulated in susceptible (UpR/DownS)

**Table S13**: Pathogen-Host Interaction (PHI-base) annotations of differentially expressed genes, with associated virulence phenotypes

**Table S14**: CAZyme annotations of differentially expressed genes

**Table S15**: Predicted effector proteins

**Table S16**: Retained intron (RI) events detected between SI and RI samples; differentially alternatively spliced genes (DAGs) are highlighted

**Table S17**: Alternative 5⍰ splice site (A5) events between SI and RI samples, with highlighted DAGs

**Table S18**: Alternative 3⍰ splice site (A3) events between SI and RI samples, with highlighted DAGs

**Table S19**: Alternative first exon (AF) events between SI and RI samples, with highlighted DAGs

**Table S20**: Skipped exon (SE) events detected between SI and RI samples, with highlighted DAGs

**Table S21**: EggNOG-based functional annotation of 22 DAGs, including ASE type, Uniprot hits, orthologs, COG categories, GO terms, EC numbers, KEGG pathways, PHI-base entries, and domain predictions

**Table S22**: PHI-base annotations of DAGs

**Table S23**: Initial set of 270 lncRNA candidates identified, with transcript IDs and lengths

**Table S24**: CPC2 results for lncRNA candidates with coding probability and predicted coding labels

**Table S25**: CPAT results for lncRNA candidates with ORF metrics and coding probabilities

**Table S26**: Final set of 150 lncRNAs with transcript lengths and coding probabilities from CPC2 and CPAT analyses

**Table S27**: Predicted subcellular localization of lncRNAs

**Table S28**: Classification of lncRNAs based on class codes and predicted lncRNA types

**Table S29**: Differentially expressed lncRNAs

**Table S30**: Final 14 DElncRNAs: 1 (highlighted one) upregulated in resistant/downregulated in susceptible (UpR/DownS) and 13 downregulated in resistant/upregulated in susceptible (DownR/UpS)

**Table S31**: Predicted cis-target genes of DElncRNAs included in the interaction network

**Table S32**: Predicted trans-target genes of DElncRNAs included in the interaction network

