## Supplementary figures and images for "Multilayered Transcriptomic Reprogramming and Spliceosomal Divergence Shape *Bipolaris sorokiniana* Pathogenicity"

### Figure S1

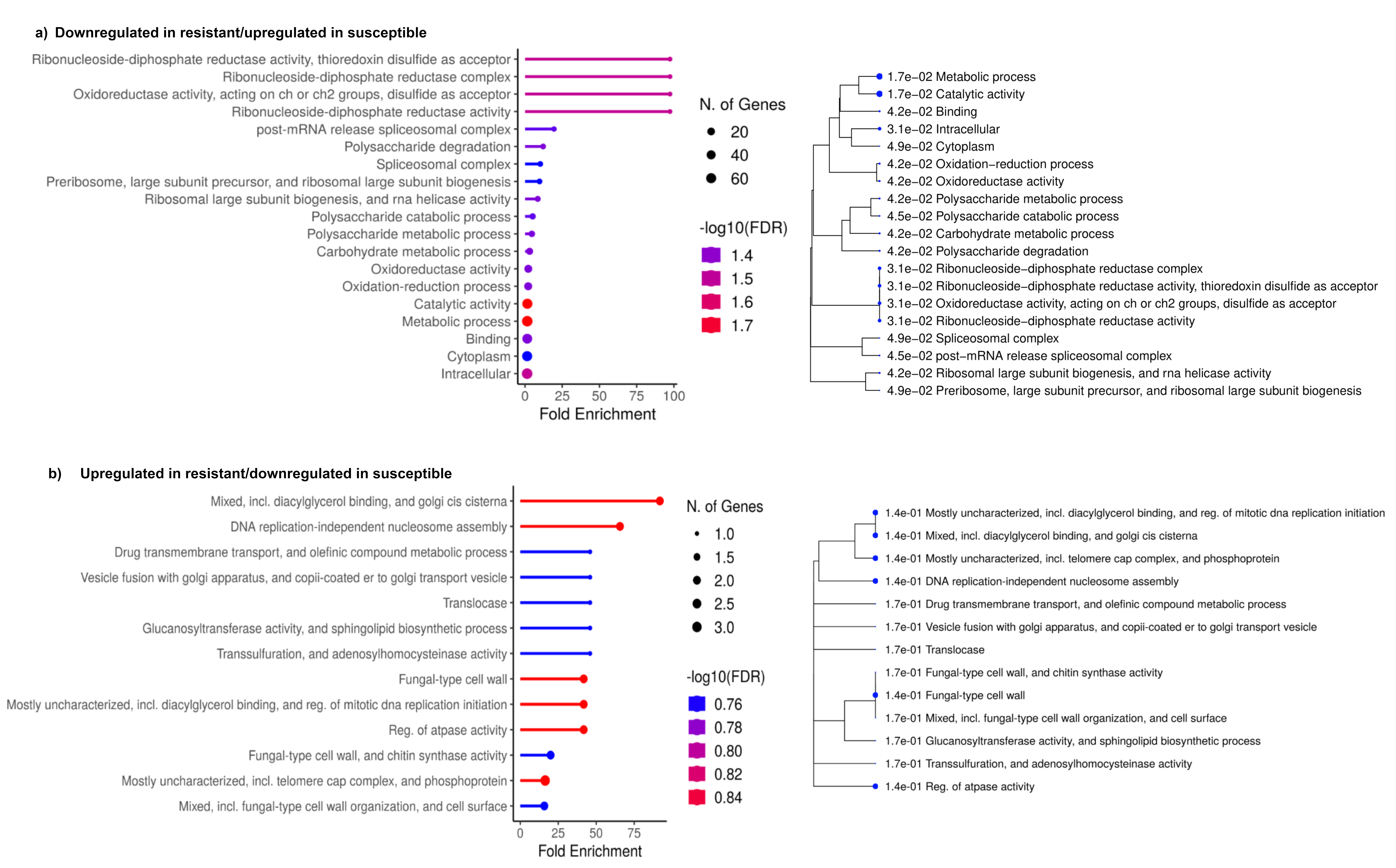

### Figure S2

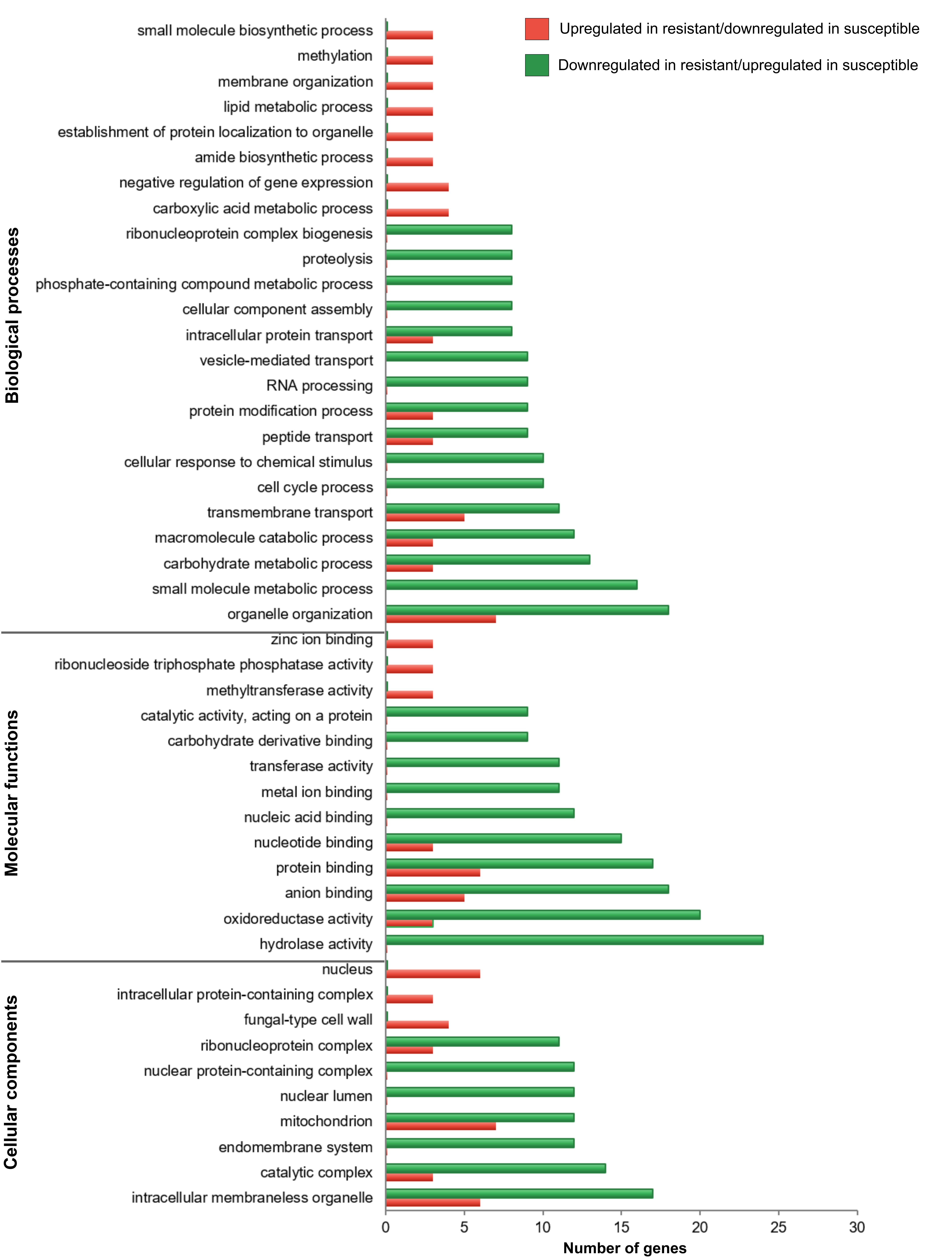

### Figure S3

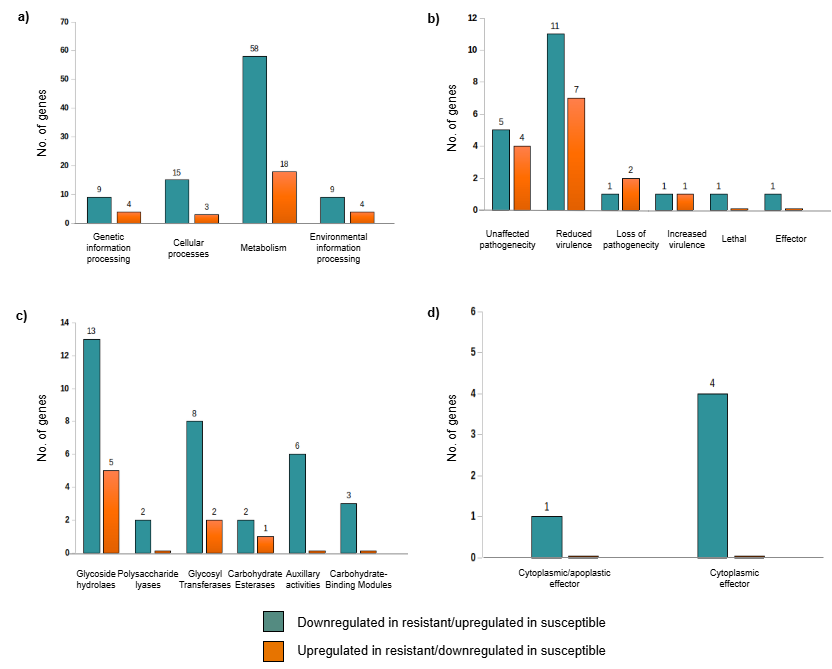

### Figure S4

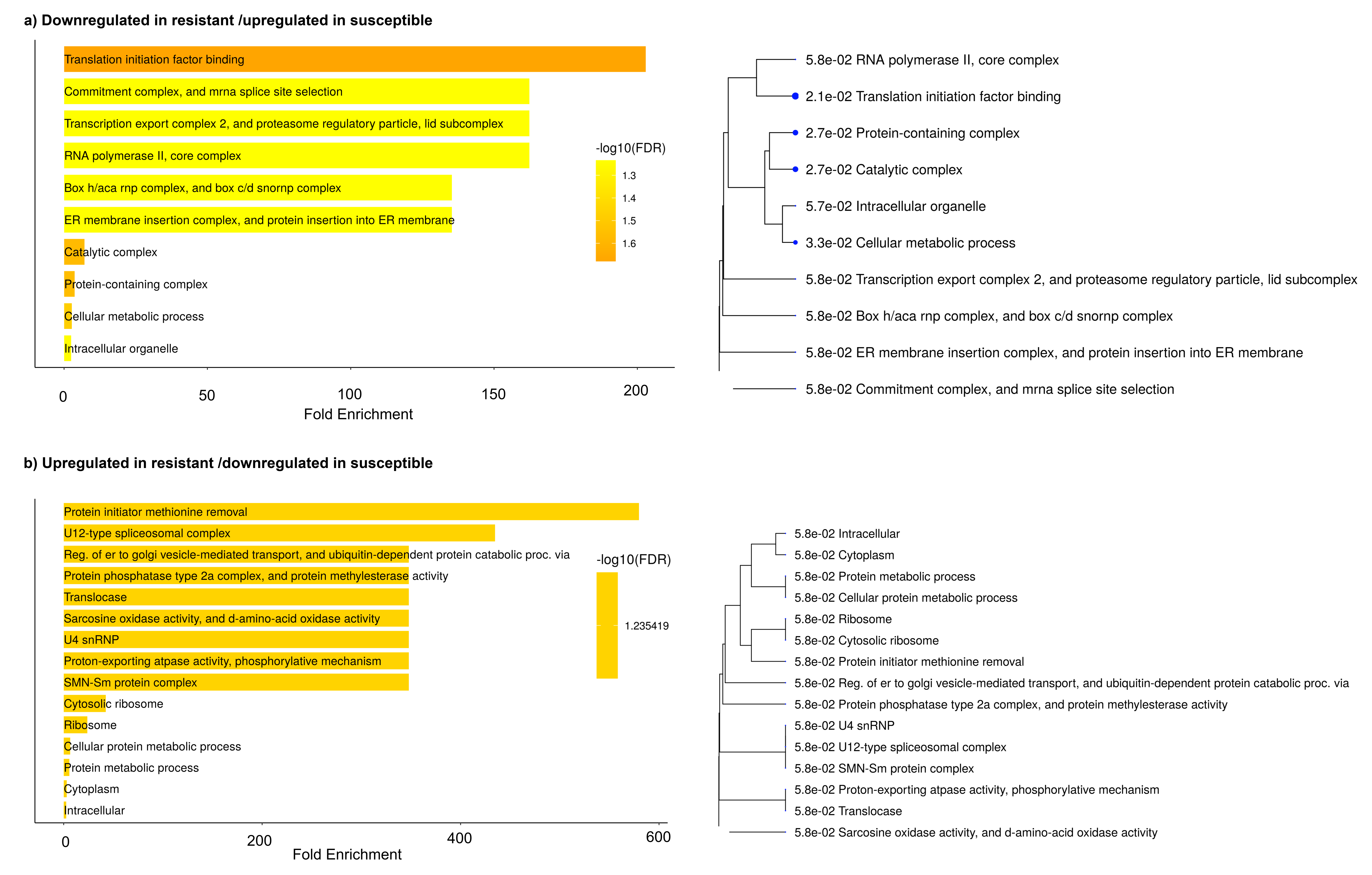

### Figure S5

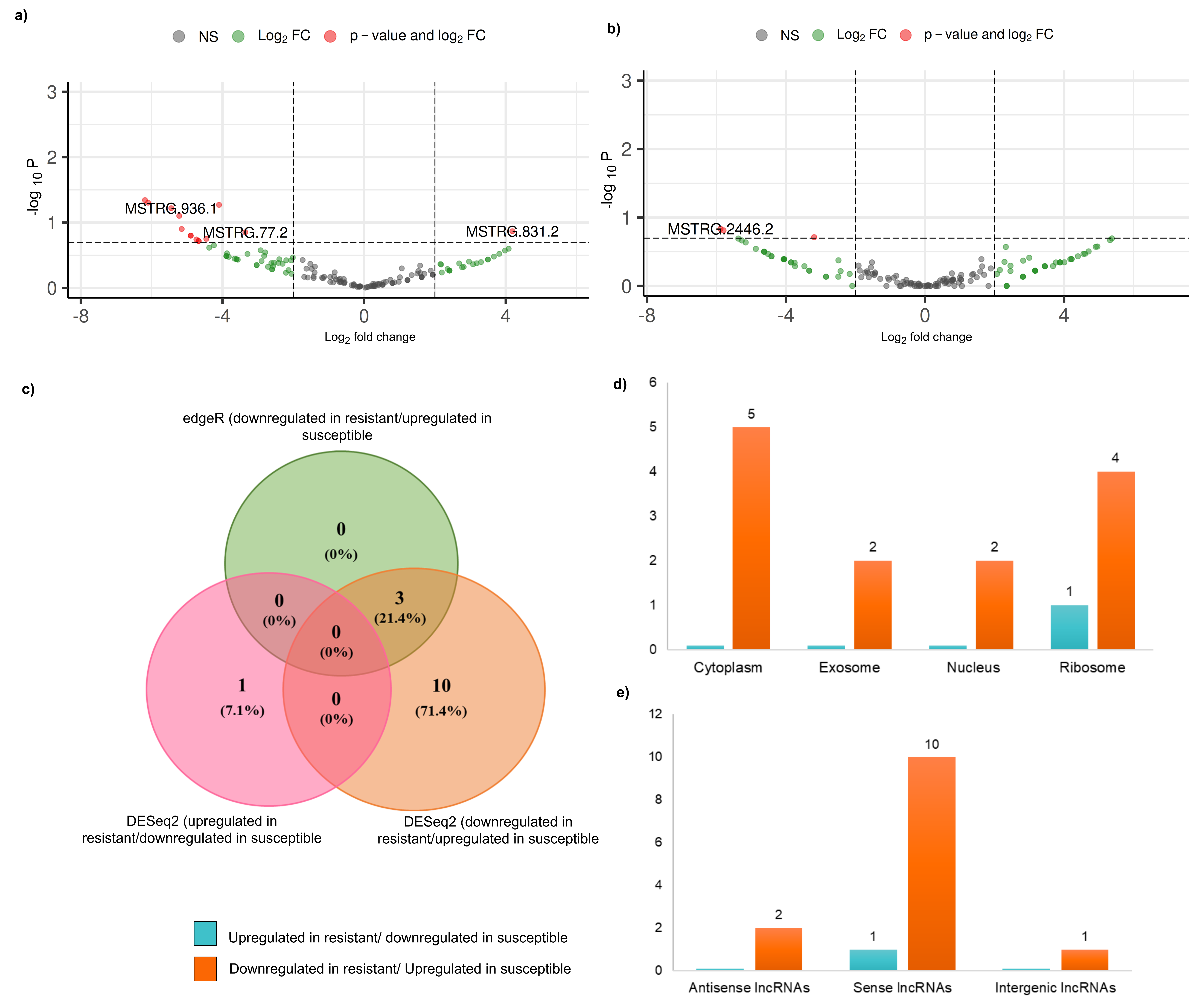

### Figure S6

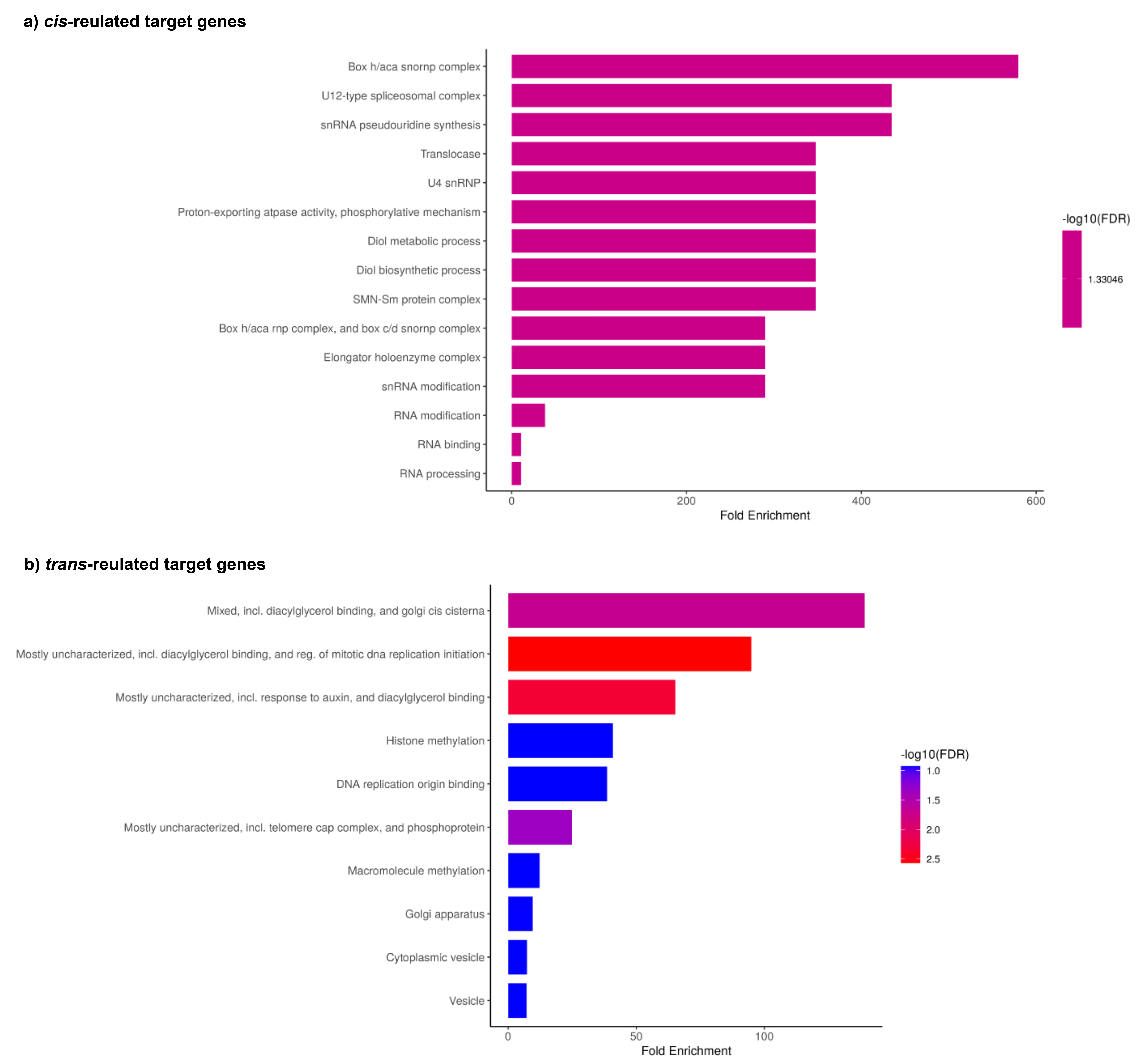
